# Deconvolution of multiplexed peptidoform mass spectra enables high-resolution profiling of complex protein modification patterns

**DOI:** 10.1101/2025.05.24.655917

**Authors:** Zhiyuan Cheng, Linhui Zhai, Li Yu, Kaifeng Chen, Wensi Zhao, Minjia Tan, Yan Fu

## Abstract

Combinatorial patterns of post-translational modifications (PTMs), like the histone code, play critical roles in regulating protein functionality. However, in bottom-up proteomics, isobaric modification peptidoforms (IMPs) frequently co-elute and co-fragment during liquid chromatography coupled with tandem mass spectrometry (LC-MS/MS), resulting in multiplexed mass spectra that cannot be properly analyzed by conventional protein identification and quantification methods. Existing solutions are either limited to highly narrow scenarios or unavailable for public use. Here, we introduce PTMdecoder, a robust and versatile software tool for discriminating and quantifying IMPs by deconvoluting their multiplexed MS/MS spectra and ion chromatograms without requiring a spectral library of individual peptidoforms. Tested on 34 samples of synthesized IMPs, PTMdecoder achieved an overall sensitivity of 95.7% for IMP identification and root mean square errors mostly <0.05 for quantification. In human histone samples, it detected and quantified 36.7–44.8% more IMPs than conventional methods, providing a higher-resolution view of histone modification patterns.

## Introduction

Post-translational modifications (PTMs) play a critical role in regulating the function and activity of proteins^1–4^. Accurate localization and quantification of PTMs are crucial for comprehensive protein characterization. In cellular contexts, certain PTMs may possess multiple potential modification sites on the protein, and these sites may be densely clustered. Importantly, the combinatorial patterns of PTMs on specific local sequences of proteins often carry significant biological implications^5–8^. For instance, histones contain numerous lysine residues that are frequently subjected to PTMs such as acetylation and methylation. The diverse PTMs on histones constitute the so-called “histone code”, which plays a key role in regulating DNA transcription^9–11^. Proteins that exhibit identical amino acid sequence but differ in their PTM patterns, encompassing distinct modification types and sites, are known as proteoforms. In bottom-up proteomics, proteoforms are enzymatically digested into peptides, yielding peptidoforms, i.e., peptides with identical sequence but different PTMs^12^. Notably, isobaric modification peptidoforms (IMPs) possess the same masses, the same or highly similar types of PTMs, and similar physicochemical properties. Therefore, they often co-elute and co-fragment during liquid chromatography coupled with tandem mass spectrometry (LC-MS/MS), generating multiplexed mass spectra^13–16^. However, current data analysis strategies typically identify only one peptide or peptidoform (the top-scoring one) per spectrum, which is subsequently utilized for quantification^17–19^. This simplified approach hinders the detection and quantification of all IMPs existing in a sample, which is increasingly critical for in-depth protein analysis, such as dynamics and crosstalk of PTMs^20,21^.

To resolve IMPs from their multiplexed spectra, several algorithms and software tools have been developed. However, they were either not designed for general scenarios or are not publicly available. Among various approaches, many were specifically tailored to particular situations, relying on knowledge derived from particular experiments or conditions. For instance, Pesavento et al.^22^ investigated the scenario of two co-existing IMPs and demonstrated that the fragment ion relative ratio (FIRR) can approximate the ratio of intact peptides. Similarly, several early-stage algorithms were designed to focus exclusively on resolving two IMPs^23–25^. The EpiProfile algorithm is capable of handling more complex mixtures (e.g., two, four, or six IMPs), but is restricted to specific histone peptides and relies on pre-established equations that are manually solved in advance^26,27^. In contrast to these specialized approaches, more general methods have also been proposed^28–30^. These methods utilize experimental fragment ions to establish linear models to resolve IMPs. Unfortunately, none of these methods have released publicly available software tools. The Iso-PeptidAce algorithm, proposed by Abishiru et al.^31^, formulates the deconvolution of multiplexed spectra as a maximum flow problem. Nevertheless, this algorithm requires individual spectra for all peptidoforms, which can be prohibitively costly or even impractical when dealing with a large number of possible peptidoforms. Moreover, the software of Iso-PeptidAce is no longer accessible, either. The aforementioned methods are primarily designed for data-dependent acquisition (DDA) mass spectra. In the data-independent acquisition (DIA) mode, multiplexed spectra are even more prevalent. However, most DIA search engines can only identify isobaric peptides with different sequences and do not discriminate isobaric peptidoforms^32–35^. A few exceptions are capable of identifying and quantifying PTM positional isomers (i.e. peptidoforms with the same modification compositions but different sites) in relatively simple patterns^36,37^. However, they need to construct a high-precision spectral library of all candidate isomers, and may meet difficulties when dealing with numerous IMPs. At present, the theoretical prediction of MS/MS spectra is largely restricted to unmodified peptides^38–40^ or those with relatively simple modification states^41–43^, and accurate spectrum prediction for arbitrarily modified peptides remains an unsolved problem. In summary, the multiplexed mass spectra of IMPs cannot yet be adequately analyzed by current proteomic software tools.

Here, we introduce PTMdecoder, a robust and versatile software tool for discriminating and quantifying IMPs in DDA-based bottom-up proteomics. Unlike existing methods, PTMdecoder imposes no limits on IMP types or numbers in multiplexed spectra and does not require a spectral library of individual peptidoforms. Instead, it directly infers IMPs from LC-MS/MS data using a two-step approach: first, it estimates the relative abundances of IMPs in each multiplexed spectrum through a fragment ion-based linear model; second, it deconvolutes the total extracted ion chromatograms (XICs) of mixtures using a non-parametric method for more accurate IMP quantification. We evaluated the performance of PTMdecoder on synthesized IMPs and biological samples for histone code analysis. On 34 samples of synthesized IMPs, PTMdecoder achieved an overall sensitivity of 95.7% for IMP identification with false positive rate of 2.4%, and root mean square errors (RMSEs) mostly <0.05 for IMP quantification. In human histone samples treated with inhibitors, PTMdecoder identified 36.7-44.8% more IMPs than conventional methods, and quantified PTM changes at levels of both individual sites and combinations. PTMdecoder aims to provide an unprecedented high resolution for PTM profiling by resolving IMPs from the multiplexed mass spectra that are widely existing but underexplored in current bottom-up proteomics.

## Results

### The overall design of PTMdecoder

Due to similar physiochemical properties, some IMPs cannot be separated during LC-MS/MS analysis, resulting in multiplexed spectra, which need to be sufficiently analyzed to uncover the present IMPs and quantify their abundances. PTMdecoder addresses this challenge by deconvoluting both the MS/MS spectra and the chromatograms, enabling accurate discrimination and quantification of IMPs coexisting in a sample (Fig. 1a).

**Figure 1.**
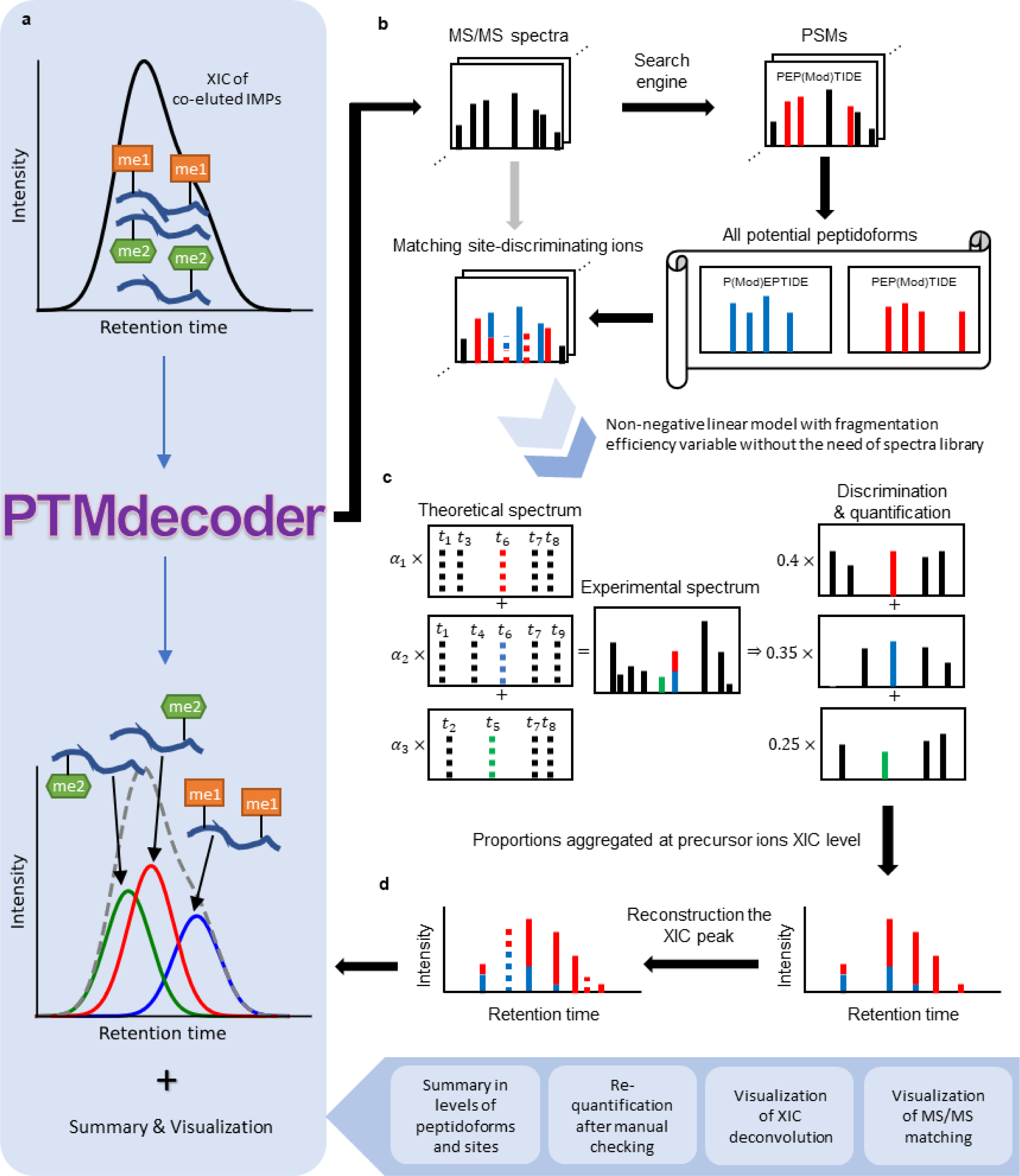
Overview of the PTMdecoder workflow. (**a**) IMPs may co-elute and co-fragment in LC-MS/MS analysis, resulting in multiplexed mass spectra. PTMdecoder deconvolutes them to discriminate and quantify individual IMPs in the mixture. (**b**) MS/MS spectra are first searched against the protein database using a conventional search engine, and the resulting peptide-spectrum matches (PSMs) with specific modifications are selected from the search results. Next, for each selected PSM, all possible IMPs are generated according to the peptide sequence, precursor mass, and considered modification types (specified by the user). Then the theoretical site-discriminating ions are calculated for each IMP and matched against the experimental spectrum. (**c**) The intensities of site-discriminating ions are used to establish a non-negative linear model, enabling the determination of the relative abundances of IMPs at MS/MS level. (**d**) Based on the relative abundances quantified at specific MS/MS acquisition time points, non-parametric methods are used to estimate the relative abundances at all time points during the elution. The total XIC of mixed precursor ions is deconvoluted into reconstructed XICs of individual IMPs. The final abundance of each IMP is computed as the area under its reconstructed XIC curve.

Given a multiplexed MS/MS spectrum and its corresponding peptide sequence, which can be identified using conventional search engines, PTMdecoder estimates the relative abundances of component IMPs through a linear model independent of individual peptidoform spectra (Methods). Specifically, it first enumerates all possible IMPs according to the peptide sequence and specified PTM types, and then utilizes the observed site-discriminating ions^44^ to detect and quantify the potential IMPs (Fig. 1b). Site-discriminating ions are those fragment ions that are informative for determining the types and positions of PTMs.

Unlike previous approaches^26–28,30^, which treat fragment ion intensities uniformly, PTMdecoder accounts for the variations in fragment ion intensities. This is because the experimental spectra typically exhibit intensity patterns that depend on the type, cleavage site, and charge state of fragment ions. To capture these patterns, PTMdecoder introduces the fragmentation efficiency variables (𝛼), and establishes a non-negative linear model based on site-discriminating ions. The model aims to minimize the difference between the observed and reconstructed intensities of these site-discriminating ions while incorporating discrimination constraints (Methods). The relative abundances of all IMPs are then derived from this model (Fig. 1c).

Subsequently, PTMdecoder quantifies IMPs through XIC deconvolution and integration. For each IMP, PTMdecoder aggregates its estimated relative abundances derived from the deconvolution of MS/MS spectra at different retention time points within the corresponding XIC peak range. Subsequently, a non-parametric method is applied to estimate the relative abundance profile and deconvolute the total XIC into multiple reconstructed component XIC curves, each for an individual IMP. Finally, PTMdecoder quantifies the IMPs by calculating the area under the reconstructed XIC curve (Fig. 1d). Only IMPs with significant relative abundance within the current XIC peak are retained for further analysis.

Additionally, PTMdecoder can quantify each PTM site by integrating the relative abundances of all IMPs containing this PTM site (Table 1), and visualize the deconvoluted XICs along with their corresponding peptidoform-spectrum matches (Supplementary Fig. 6-7). For downstream data analysis, PTMdecoder supports the comparison of quantifications across different runs, the quantifications of normalization peptides (used to normalize the changes of ratio in quantification between runs), and re-quantification based on manual alignment of XIC peaks across runs.

**Table 1.** The normalized ratio (SAHA/DMSO) of IMPs with acetylation on H4K12 in two replicates.

| Sequence | Charge | Normalized ratio<br>(rep 1) | Normalized ratio<br>(rep 2) |
| --- | --- | --- | --- |
| _GGK{Acetyl}GLG**K**{Acetyl}GGAK_ | +1 | 2.26 | 2.27 |
| _GGK{Acetyl}GLG**K**{Acetyl}GGAK_ | +2 | 2.61 | 2.31 |
| _GGK{Acetyl}GLG**K**{Acetyl}GGAK{Acetyl}R_ | +2 | 10.33 | 8.51 |
| _GGK{Acetyl}GLG**K**{Acetyl}GGAK{Acetyl}R_ | +3 | 10.59 | 7.05 |
| _GKGGK{Acetyl}GLG**K**{Acetyl}GGAK_ | +2 | 3.67 | 2.19 |
| _GKGGK{Acetyl}GLG**K**{Acetyl}GGAK_ | +3 | 4.01 | 2.04 |
| _GKGGK{Acetyl}GLG**K**{Acetyl}GGAK{Acetyl}R_ | +2 | 11.03 | 6.07 |
| _GKGGK{Acetyl}GLG**K**{Acetyl}GGAK{Acetyl}R_ | +3 | 10.07 | - |
| _GK{Acetyl}GGKGLG**K**{Acetyl}GGAK_ | +3 | 3.38 | 2.00 |
| _GK{Acetyl}GGKGLG**K**{Acetyl}GGAK{Acetyl}R_ | +2 | 16.80 | 7.01 |
| _GK{Acetyl}GGKGLG**K**{Acetyl}GGAK{Acetyl}R_ | +3 | 11.99 | - |
| _GK{Acetyl}GGK{Acetyl}GLG**K**{Acetyl}GGAK_ | +2 | 6.76 | 6.15 |
| _GK{Acetyl}GGK{Acetyl}GLG**K**{Acetyl}GGAK{Acetyl}R_ | +2 | 24.90 | 22.35 |
| _GK{Acetyl}GGK{Acetyl}GLG**K**{Acetyl}GGAK{Acetyl}R_ | +3 | 23.48 | 18.81 |
| _GLG**K**{Acetyl}GGAKR_ | +2 | 0.99 | 2.10 |
| _GLG**K**{Acetyl}GGAK_ | +1 | 1.44 | 1.53 |
| _GLG**K**{Acetyl}GGAK_ | +2 | 1.25 | 1.29 |
| _GLG**K**{Acetyl}GGAK{Acetyl}R_ | +1 | 5.69 | 5.43 |
| _GLG**K**{Acetyl}GGAK{Acetyl}R_ | +2 | 4.29 | 4.08 |

### Performance evaluation of PTMdecoder using synthesized peptides

Sixteen peptides with distinct IMPs were synthesized to validate the performance of PTMdecoder (Fig. 2a, Methods). These IMPs were generated by incorporating three types of PTMs—acetylation, propionylation, and propionylation-methylation on lysine—into the human histone H4 4-17 peptide in 16 unique combinations (Fig. 2a). Specifically, we synthesized 16 peptides with identical sequences but harboring different acetylation and methylation patterns at various lysine sites. Subsequently, propionylation was performed at the peptide level to label all free amino groups, including the N-terminal amino group, unmodified lysine side chains, and mono-methylated lysine residues. Through this in vitro propionylation-based chemical derivatization, we obtained 16 IMPs with identical molecular weights but distinct combinations of different modification types. This design not only produced positional isomers but also yielded peptidoforms with different types of modifications. Initially, we analyzed the individual-component samples of the 16 IMPs separately. Then, a total of 18 mixed samples were prepared by combining the 16 synthesized IMPs in specific proportions (parts) as shown in Fig. 2b. These 18 mixed samples were organized into 9 pairs, with each pair containing the same set of IMPs, one in equal proportions and the other in unequal proportions. All the 34 samples were analyzed by LC-MS/MS (Methods). The resulting MS/MS data showed that the IMPs in all mixed samples partially or even completely co-eluted and co-fragmented. Most mixed samples (including the two containing all 16 IMPs) produced single XIC peaks across the entire chromatographic gradient (Supplementary Fig. 2). Such serious co-elution would pose a significant challenge to conventional peptide identification methods.

**Figure 2.**
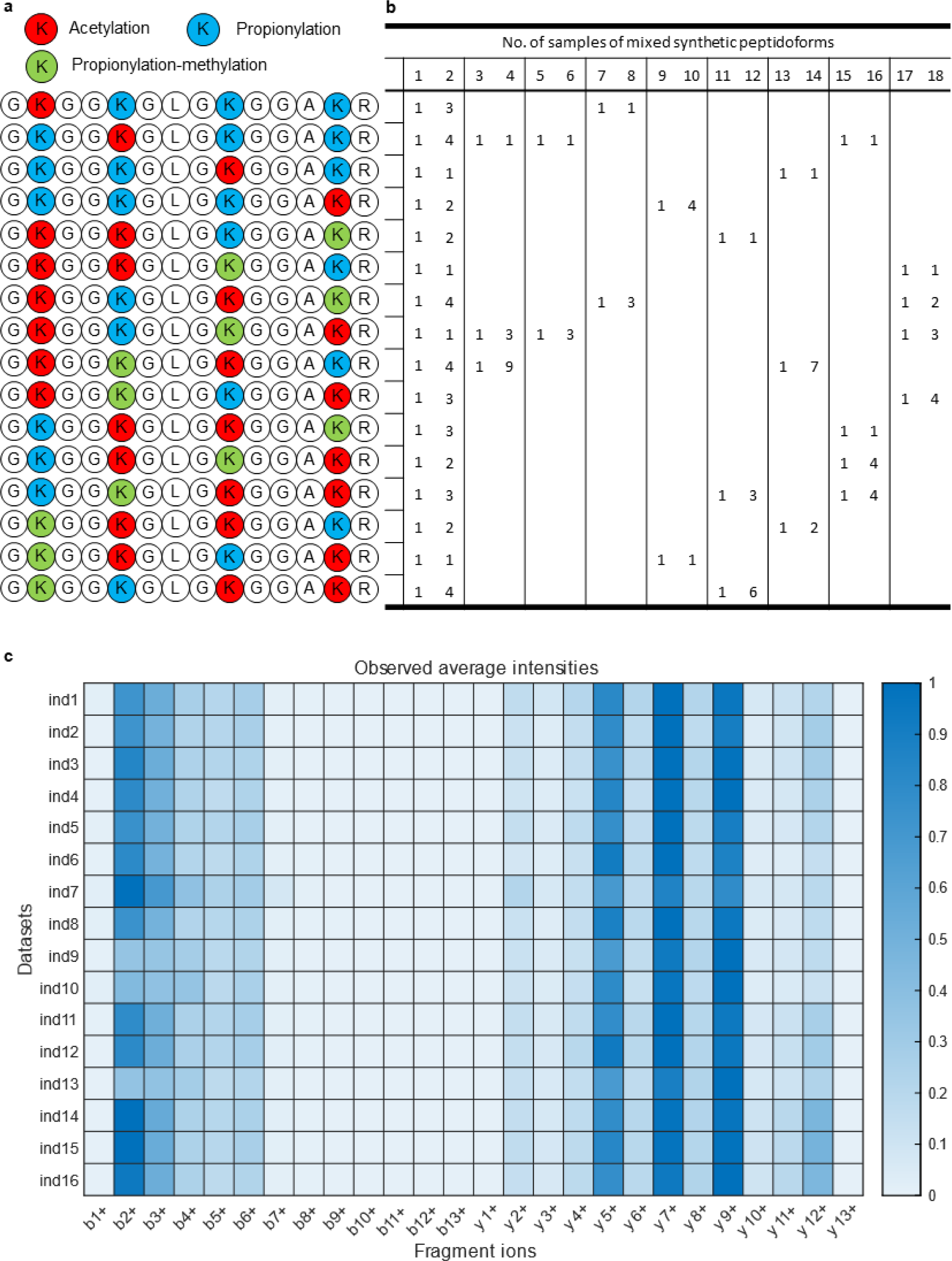
Design of synthesized IMP samples and analysis of fragmentation efficiencies. (**a**) Synthesized human histone H4 4-17 peptides with various PTMs. Sixteen IMPs were synthesized, which have identical precursor mass but different modification compositions or sites. The modifications included acetylations, propionylations, and propionylation-methylations on the arginine residues, and propionylation on the peptide N-terminus (not shown). (**b**) Design matrix of mixed samples. A total of 18 mixed samples were generated in different proportions (parts). Each row corresponds to an IMP in (a), and each column corresponds to a mixed sample. (**c**) Heatmap of fragmentation efficiencies (observed average intensities of fragment ions). The fragmentation efficiencies were different between fragment ion classes but similar between IMPs. Each row corresponds to an individual IMP sample.

We first validated the assumption that IMPs share similar fragmentation efficiencies using the 16 single-IMP samples. The fragment ions were categorized into classes based on their terminal type, number of residues, and charge state, with each class assigned a corresponding efficiency variable. The real fragmentation efficiencies (fragment ion intensities) were calculated for all 16 IMPs. In Fig. 2c, each row represents the average intensities of fragment ions derived from a synthesized IMP in Fig. 2a. Despite the different modifications present in the fragment ions, the average intensities of fragment ions in the same class are remarkably consistent across IMPs, whereas significant differences were observed between classes. As shown in Supplementary Fig. 3, the fragmentation efficiencies estimated by the variable model of PTMdecoder closely align with the observed intensities as shown in Fig. 2c. A similar trend in the fragmentation efficiencies was also observed in the 18 mixed samples, as depicted in Supplementary Fig. 4. All these results demonstrate the rationality of our variable model of fragmentation efficiency, which enables PTMdecoder to perform library-free analysis of IMPs.

Then we detected coeluted IMPs by deconvoluting the multiplexed mass spectra using PTMdecoder. Since the exact proportions of the synthesized IMPs in mixtures were predetermined during sample preparation, it is straightforward to validate the presence of IMPs discriminated by PTMdecoder and assess the quantification accuracy of the percentage of each IMP in the mixture. The IMPs identified in a sample can be divided into two groups: true positives, which were indeed added to the mixture, and false positives, which were not present in the mixture but were incorrectly identified. For the coeluted IMPs detected at the MS/MS spectrum level, a filtering threshold was applied to exclude potentially unreliable IMPs that had very low relative abundances. Specifically, IMPs with abundances less than the maximum abundance of co-eluted IMPs multiplied by a threshold factor were removed. The true positive rate (TPR) and false positive rate (FPR) across varying filtering thresholds on the 34 datasets (16 individual IMP samples and 18 mixed samples) were calculated and formed the ROC curves (Fig. 3a). The AUC was calculated to characterize the discriminating capacity of PTMdecoder, yielding a value of 0.9969. To balance the TPR and FPR, a filtering threshold of 0.1 was uniformly chosen for all datasets, as highlighted by the red circle in Fig. 3a. Utilizing this threshold, the variable model of fragmentation efficiency in PTMdecoder exhibited excellent performance across these 34 datasets, as illustrated in Fig. 3b, achieving a recall of 100% on 31 out of 34 datasets while maintaining a low overall FPR of 2.4%. On the four datasets with false positives identified, we observed multicollinearity among the matched ions derived from identified IMPs. The discrimination and quantification results for the 9 unequally mixed samples are shown in Fig. 3c as an example. In the figure, distinct IMPs are represented by different colors, while the segment lengths reflect their relative abundances. The estimated relative abundances are highly consistent with the real ratios in the mixed samples. In the ‘mix1’ sample consisting of 16 IMPs, PTMdecoder successfully detected as many as 13 co-eluted IMPs from a single MS/MS spectrum, revealing the remarkable co-elution phenomenon. Combining the results of all identified MS/MS spectra, all 16 IMPs were detected.

**Figure 3.**
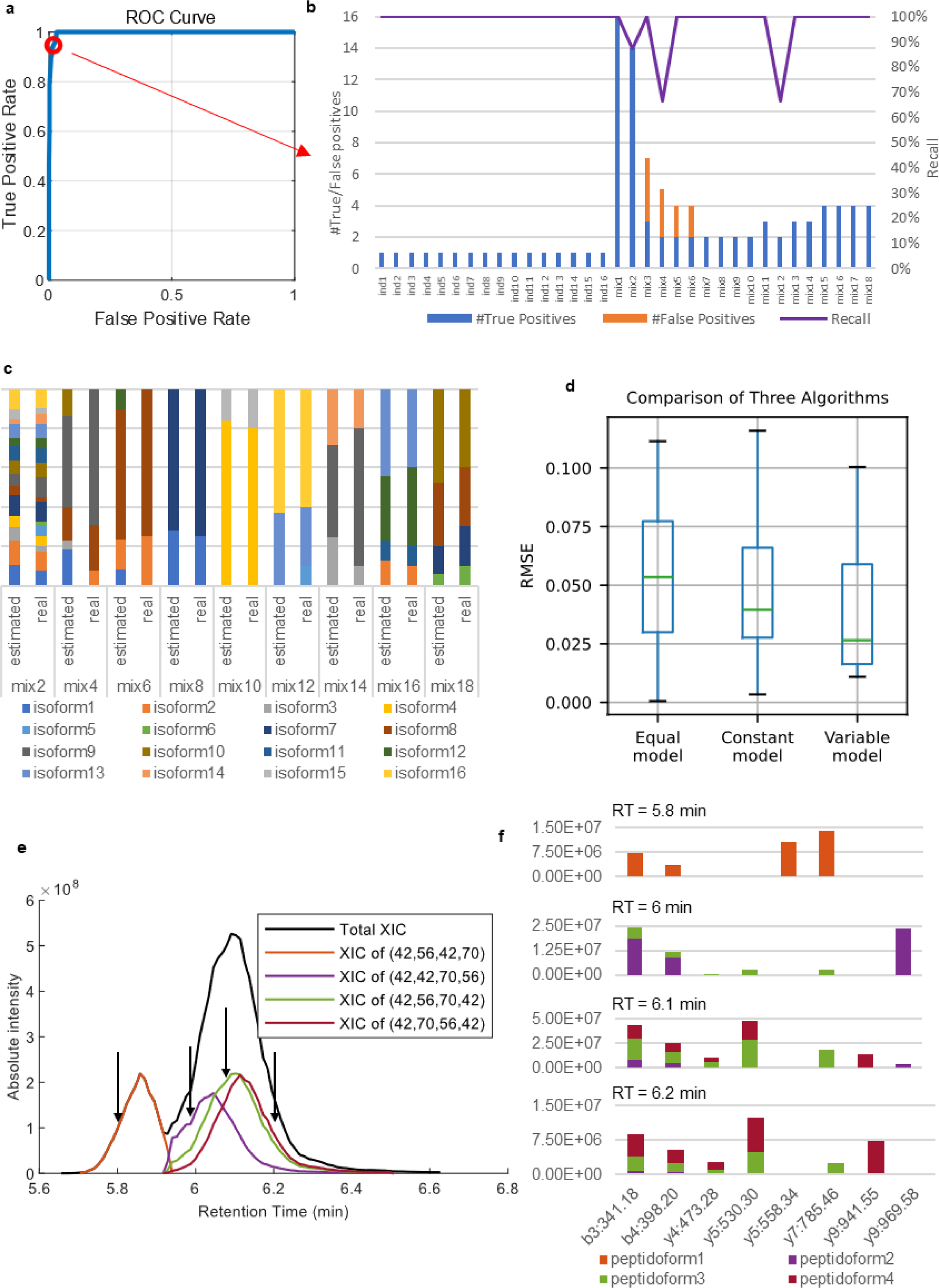
Discrimination and quantification of synthesized IMPs. (**a**) The overall receiver operating characteristic (ROC) curve for the 34 samples of synthesized IMPs. (**b**) Numbers of true/false positives (left) and recall (right) for each of the 34 samples, obtained using PTMdecoder’s fragmentation efficiency variable model with the threshold denoted by the red circle in (a). (**c**) Comparison between estimated and real mixing proportions of IMPs across nine unequally-mixed samples. (**d**) Box plot of quantification RMSEs for three fragmentation efficiency models across 18 mixed samples. The ‘Equal model’ assumes identical fragmentation efficiencies for all fragment ion classes; the ‘Constant model’ assigns distinct, pre-estimated constant efficiencies to each class; the ‘Variable model’ assumes different fragmentation efficiencies between ion classes and estimates them simultaneously with the quantification model. It is a novel model introduced in this work and used by PTMdecoder. (**e**) An example illustrating the deconvolution of an XIC peak for mixture into individual IMP components. The four vectors in the legend represent four IMPs in Fig. 2a, with the numbers indicating modification types (nominal masses) on four arginine residues, 42 for acetylation, 56 for propionylation, and 70 for propionylation-methylation. (**f**) Deconvoluted intensities of selected fragment ions across four MS/MS spectra during the elution profile shown in (e).

In addition to treating fragmentation efficiencies as variables (our proposed variable model), we implemented two established approaches for comparative analysis: the equal model and the constant model^28,29^ (Supplementary Fig. 1). Note that no suitable software tools are currently available for these models. The equal model operates under the assumption of equal fragmentation efficiencies across all fragment ion types. It represents a class of methods that directly utilize the FIRR to quantify each IMP in multiplexed MS/MS spectra^22,28^. In contrast, the constant model treats fragmentation efficiencies as constants related to the class of fragment ions. However, unlike the variable model, the constant model estimates fragmentation efficiencies before the deconvolution of multiplexed MS/MS spectra^29^. On the synthesized-peptide samples, the variable model outperformed the other two models in terms of the numbers of true/false positives, the abundances of false positives, and the quantification RMSE (root mean squared error) (Fig. 3d, Supplementary Fig. 5).

The deconvolutions of XICs of all 18 mixed samples are presented in Supplementary Fig. 6. An example of an XIC peak deconvoluted by PTMdecoder is shown in Fig. 3e-f. This XIC peak originated from an equally-mixed sample of four IMPs, as shown in the legend. Notably, some of the IMP peptides could be partially separated by reversed HPLC. In contrast to the methods that rely exclusively on unique fragment ions, PTMdecoder resolves the differential contributions of the shared fragment ions from different IMPs. Fig. 3f shows the deconvolution of the intensities of some site-discriminating ions in MS/MS spectra at the four retention times marked by black arrows in Fig. 3e. The matching fragment peaks of top-scoring and all PTMdecoder-discriminated IMPs at these four retention times (Supplementary Fig. 7) demonstrate that these spectra were indeed generated from multiple IMPs. The relative abundances of IMPs at each time point within the retention time range were estimated using a non-parametric method. Subsequently, the total XIC peak in Fig. 3e was deconvoluted into four reconstructed XIC peaks, each corresponding to one of the four IMPs. The total XIC of the mixture is shown by the black line, while the reconstructed XIC curves of four composing IMPs are depicted in color. Although the deconvolution of MS/MS spectra at each retention time yielded clearly unequal proportions of the four IMPs, the areas under the reconstructed XIC peaks of the IMPs are pretty close to each other, indicating that the quantification by PTMdecoder is reliable. In Fig. 3e, the peptidoform “(42, 56, 70, 42)”, i.e., “_{Propionyl}GK{Acetyl}GGK{Propionyl}GLGK{Propionyl-methyl}GGAK{Acetyl}R_”, lacks unique site-discriminating ions, since all its fragment ions can also be produced by other existing IMPs. Nevertheless, PTMdecoder successfully detected and quantified this peptidoform. PTMdecoder also accurately filtered out the 12 IMPs that were not included in this mixture. This case demonstrates PTMdecoder’s capability in resolving co-eluted IMPs even when no unique site-discriminating ions exist.

### Application to the identification of upregulated histone methylation sites in MCF7 cell samples treated with JIB-04

To assess the performance of PTMdecoder on complex biological samples, we applied it to identify and quantify histone methylation changes in MCF7 cells with JIB-04 treatment, a histone demethylase pan-inhibitor. Compared with the DMSO-treated control group, histone methylation levels in the JIB-04-treated group are expected to be increased.

MCF7 cells were cultured and subsequently divided into two treatment groups (DMSO and JIB-04), with each group containing two biological replicates (Fig. 4a). Histones were extracted, separated by SDS-PAGE, and stained with Coomassie Brilliant Blue (Fig. 4b). The histone bands were excised, subjected to in-gel tryptic digestion, and analyzed by LC-MS/MS. The resulting mass spectra were analyzed by using Mascot search engine, and the Mascot identification results were analyzed using PTMdecoder (Methods). To achieve more accurate quantification at both the peptidoform and PTM site levels, we manually performed XIC peak matching across runs with the assistance of Xcalibur Qual Browser. Subsequently, the IMPs were then re-quantified within the aligned retention time range using PTMdecoder.

**Figure 4.**
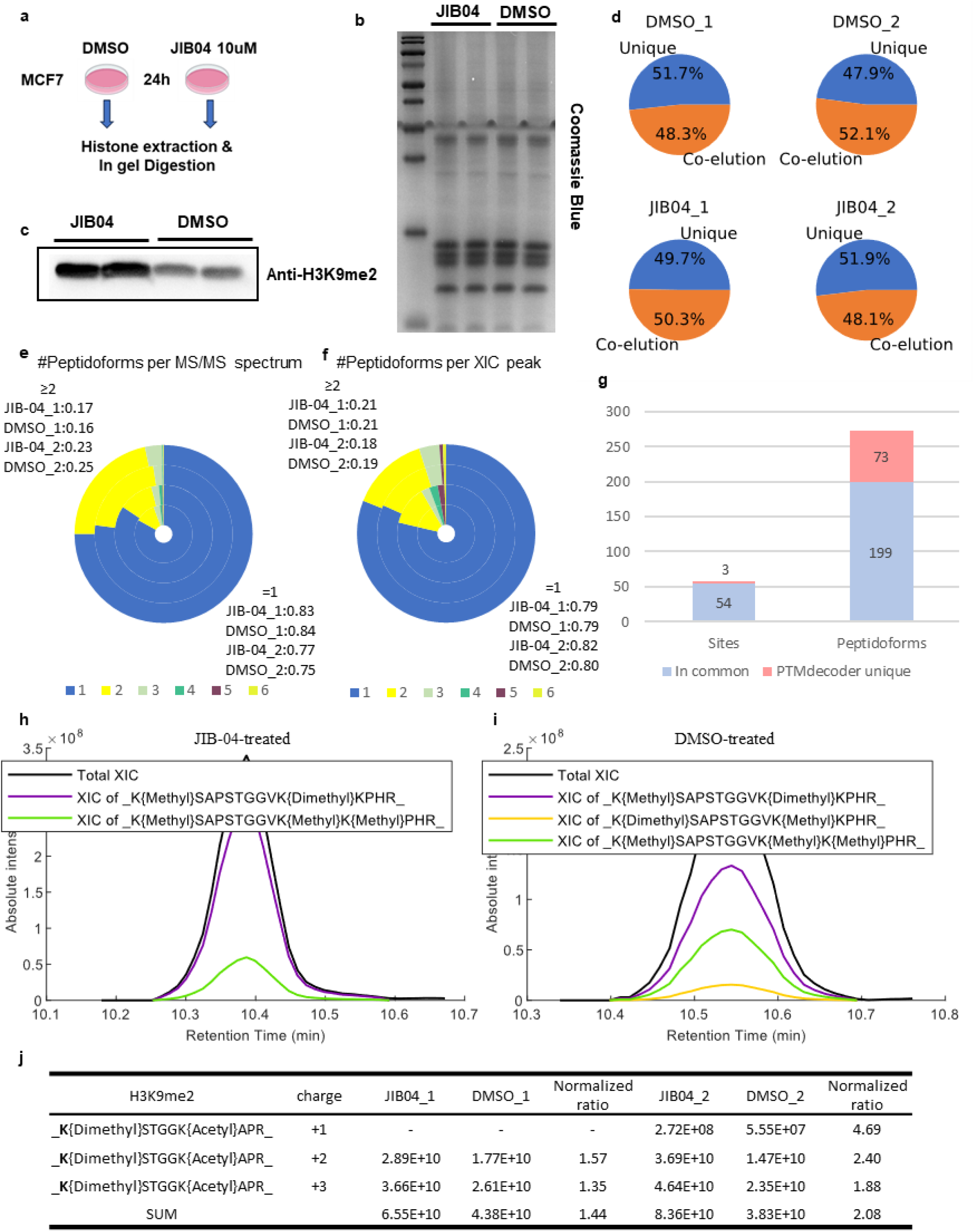
Analysis of histone methylation changes in MCF7 cells treated with JIB-04 or DMSO by PTMdecoder. (**a**) Experimental design. MCF7 cells were treated with either DMSO or JIB-04, followed by histone extraction and in-gel digestion. (**b**) SDS-PAGE separation of extracted histones. (**c**) WB analysis of histone H3K9me2 of JIB-04-treated versus DMSO-treated MCF7 cells. (**d**) Proportions of co-eluted and uniquely eluted IMPs in four runs. (**e**) Distribution of the number of IMPs identified per MS/MS spectrum across four datasets: JIB-04_1 (innermost), DMSO_1, JIB-04_2, and DMSO_2 (outermost). (**f**) Distribution of the number of IMPs identified per XIC peak across the same four datasets in (e). (**g**) Increases in the numbers of modification sites and IMPs identified by PTMdecoder compared with the conventional approach to peptide identification. (**h**) and (**i**) Deconvolutions of an XIC peak from the JIB-04-treated sample JIB-04_2 and the corresponding XIC peak from the DMSO-treated sample DMSO_2. (**j**) Quantified abundances and normalized ratios (JIB-04/DMSO) of one IMP in three charge states corresponding to H3K9me2.

PTMdecoder consistently revealed the co-elution of IMPs as a common phenomenon across multiple analytical levels. In all four LC-MS/MS runs, approximately 50% of IMPs exhibited retention time overlaps with other IMPs (Fig. 4d). At the MS/MS spectrum level, after excluding IMPs outside the aligned retention time ranges, 15% to 25% of the spectra were recognized as multiplexed spectra containing two or more IMPs (Fig. 4e). At the XIC peak level, approximately 20% of the XIC peaks corresponded to mixed IMPs (Fig. 4f).

The significance of deconvoluting multiplexed MS/MS spectra was further validated on these datasets. To demonstrate this, we implemented an alternative workflow (referred to as “top-1 mode”) in which only the top-ranked peptidoform from the search engine’s identification results was assigned to each MS/MS spectrum during the deconvolution step. Subsequently, the data were processed through PTMdecoder’s standard workflow for mixed XIC peak deconvolution and IMP quantification, followed by manual retention time alignment and re-quantification. Comparative analysis revealed that the original PTMdecoder workflow outperformed the top-1 mode, identifying 73 more IMPs (36.7% increase) and 3 additional PTM sites (5.6% increase) (Fig. 4g).

Moreover, PTMdecoder quantified subtle differences in change ratios between co-eluted IMPs in response to different treatments. For example, Fig. 4h and 4i show a pair of deconvoluted XIC peaks matched between JIB-04-treated and DMSO-treated samples. PTMdecoder identified two common IMPs, “_K{Methyl}SAPSTGGVK{Dimethyl}KPHR_” and “_K{Methyl}SAPSTGGVK{Methyl}K{Methyl}PHR_”, in the two treatments, with quantification change ratios of 1.82 and 0.74, respectively. With the top-1 mode, only the former peptide was identified, and its ratio of changes in quantification was 1.34. The evidence of fragment ion peaks for these IMPs is presented in Supplementary Fig. 8.

For H3K9me2 (dimethylation in histone H3 at position K9), PTMdecoder identified one peptidoform in three charge states, as shown in Fig. 4j. The ratios of changes between JIB-04-treated and DMSO-treated cells were similar across the two replicates. The ratios of change at the peptidoform level were consistent with the site level, i.e., SUM in Fig. 4j. These results collectively indicated that the JIB-04 treatment indeed promoted the H3K9me2 upregulation. This observation was further validated by Western blotting (WB) analysis (Fig. 4c). Since only this peptidoform with three charges associated with H3K9me2 was identified in these datasets, the WB validation also demonstrated the accuracy of the quantification of PTMdecoder at the peptidoform level.

### Application to histone acetylation analysis in MCF7 cell samples treated with SAHA

We further employed PTMdecoder to analyze acetylation patterns in MCF7 cells treated with the histone deacetylase inhibitor SAHA. MCF7 cells were cultured and treated with SAHA or DMSO in two biological replicates, following the same procedure as in the JIB-04 experiment. (Fig. 5a-b).

**Figure 5.**
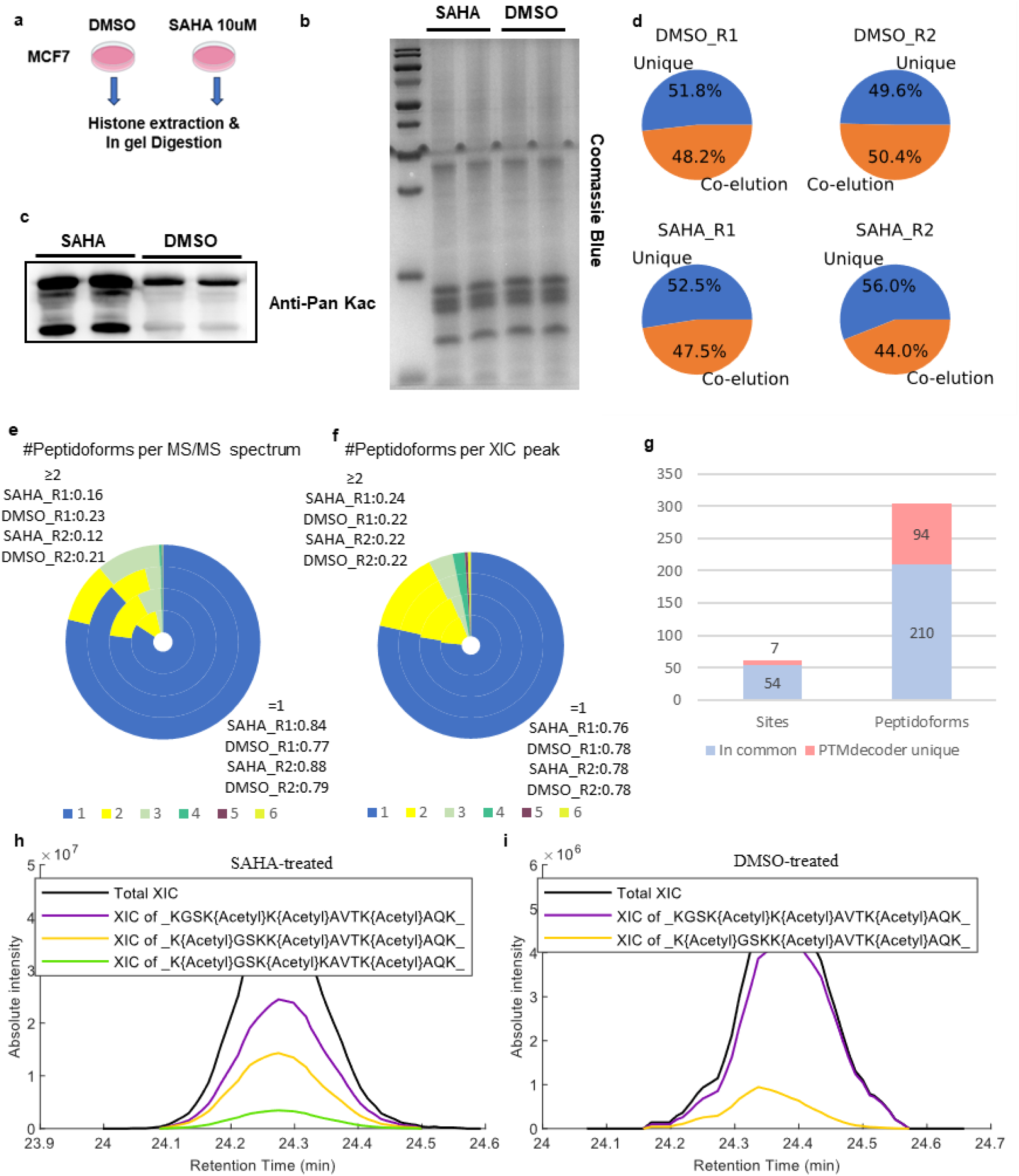
Analysis of histone acetylation changes in MCF7 cells treated with SAHA or DMSO by PTMdecoder. (**a**) Experimental design. MCF7 cells were treated with either DMSO or SAHA, followed by histone extraction and in-gel digestion. (**b**) SDS-PAGE separation of extracted histones. (**c**) WB analysis of histone pan-acetylation of SAHA-treated versus DMSO-treated MCF7 cells. (**d**) Proportions of co-eluted and uniquely eluted IMPs in four runs. (**e**) Distribution of the number of IMPs identified per MS/MS spectrum across four datasets: SAHA_R1 (innermost), DMSO_R1, SAHA_R2, and DMSO_R2 (outermost). (**f**) Distribution of the number of IMPs identified per XIC peak across the same four datasets in (e). (**g**) Increases in the numbers of modification sites and IMPs identified by PTMdecoder compared with the conventional approach to peptide identification. (**h**) and (**i**) Deconvolutions of an XIC peak from the SAHA-treated sample SAHA_R2 and the corresponding XIC peak from the DMSO-treated sample DMSO_R2.

PTMdecoder also revealed that over 50% of the IMPs co-eluted with others across all four runs (Fig. 5d). After excluding IMPs outside the aligned retention time range, PTMdecoder found that the proportions of multiplexed MS/MS spectra and XICs of mixed IMPs in the SAHA dataset were similar to that in the JIB-04 dataset (Fig. 5e-f). Comparative analysis revealed that the original PTMdecoder workflow outperformed the top-1 mode, identifying 94 additional IMPs (44.8% increase) and 7 additional PTM sites (13.0% increase) (Fig. 4g). As an example, Fig. 5h and 5i illustrate a pair of deconvoluted XIC peaks matched between SAHA-treated and DMSO-treated samples. PTMdecoder identified two common IMPs, “_KGSK{Acetyl}K{Acetyl}AVTK{Acetyl}AQK_” and “_K{Acetyl}GSKK{Acetyl}AVTK{Acetyl}AQK_”, in the two treatments, with quantification change ratios of 5.48 and 17.68, respectively. In contrast, with the top-1 mode, only the former peptide was identified, and its quantification change ratio was 8.0108. The evidence of fragment ion peaks for these IMPs is presented in Supplementary Fig. 9.

WB analysis confirmed the upregulation of histone acetylation in SAHA-treated samples (Fig. 5c). PTMdecoder identified 30 acetylation sites observed in both SAHA and DMSO-treated samples. Visualization of ratios of changes in the quantification of these sites was shown in Supplementary Fig. 10-11. Each acetylation site was supported by several peptidoforms. For example, Table 1 shows the IMPs supporting H4K12ac (acetylation at K12 of histone H4). This indicated that the ratios of changes between SAHA and DMSO-treated samples in the quantification of the same peptidoforms across different charge states were consistent, while the ratio of changes of different peptidoforms may differ. This pattern was consistent in both replicates, highlighting the capability and stability of PTMdecoder in quantifying the IMPs and providing detailed comparisons between treatment and control at both the PTM site and peptidoform levels.

In conclusion, compared to traditional methods, PTMdecoder showed much higher sensitivity in the identification of histone PTM patterns through efficient deconvolution of multiplexed spectra. Furthermore, it also showed powerful capability in precisely quantifying PTM combinations in addition to individual PTM sites and effectively screening for valuable regulatory sites associated with modification differences.

## Discussion

In LC-MS/MS-based bottom-up proteomics, co-eluted and co-fragmented IMPs bring huge difficulties to the comprehensive characterization of combinatorial PTM patterns. In this study, we proposed PTMdecoder, a software tool to discriminate and quantify co-eluted IMPs without the need for a spectral library. This addresses a significant challenge in the field, as existing approaches either lack dedicated software implementations^29–31^ or are constrained in handling diverse numbers and types of IMPs^25–27^.

We demonstrated that the linear model we used is highly effective for deconvoluting multiplexed MS/MS spectra because, ideally, the experimental spectra are generated by summing up the individual spectra of composing IMPs. Although previous studies have successfully utilized the ratio of fragment ion abundances to approximate IMP abundances^22,28–30^, our work highlights the critical role of fragmentation efficiency in library-free deconvolution of multiplexed spectra. PTMdecoder quantifies the IMPs by deconvoluting XICs of co-eluted IMPs based on the deconvolution results of multiplexed MS/MS spectra. This approach is more reasonable than the simplified quantification methods, such as spectral count or summed precursor intensities of identified peptides.

Systematic evaluation of PTMdecoder was performed on both synthesized peptides and biological samples. Its accuracy in resolving IMPs was rigorously validated using a diverse set of synthesized peptides. Its applicability for histone modification analysis was demonstrated on MCF7 cells treated with JIB-04 or SAHA. The results reveal the prevalence of IMP co-elution and underscore the critical importance of deconvoluting multiplexed mass spectra to avoid missing significant IMPs in downstream analyses.

While PTMdecoder exhibits strong performance, it still has certain limitations. First, during the deconvolution of MS/MS spectra, multicollinearity among matched fragment ions derived from IMPs may lead to the occurrence of false positives. While this does not affect site-level quantification accuracy, it may introduce errors at the peptidoform level. Second, PTMdecoder currently requires manual XIC alignment between runs before re-quantification. In controlled experiments, proper XIC alignment can reduce false positives and yield more reasonable XIC deconvolution, thereby enhancing the accuracy of peptide quantification. In the future, we will focus on overcoming the above limitations. We also intend to handle the DIA datasets containing co-eluted IMPs. By utilizing the elution profiles of site-discriminating ions, we aim to better eliminate the noise interference and achieve more accurate deconvolution. In addition, although PTMdecoder is designed for bottom-up proteomics, we anticipate that it is adaptable to middle-down and top-down proteomics.

In the life science research, high-precision analysis and quantitative characterization of PTM sites have emerged as a core technical approach for deciphering regulatory networks within living systems. PTMdecoder enables high-resolution dynamic monitoring of PTM sites and patterns, and tracking changes in modification abundance under varying physiological and pathological conditions. It can be potentially used to screen for modification sites with significant biological relevance, providing robust research targets and reliable data support for elucidating dynamic modification regulatory mechanisms and uncovering disease-associated molecular events.

## Methods

### Synthesized peptide sample processing

Sixteen different IMPs of human H4 peptide 4-GKGGKGLGKGGAKR-17, each containing distinct lysine sites with acetylation or methylation modifications, were commercially synthesized (Fig. 2a). Subsequently, propionylation was performed at the peptide level to label all free amino groups, including the N-terminal amino group, unmodified lysine side chains, and mono-methylated lysine residues. Finally, the sixteen samples of single IMPs and eighteen mixed samples comprising multiple IMPs in different proportions were prepared. The proportions (parts) of IMPs in each sample are shown in Fig. 2b. The IMPs peptide was resuspended in Buffer A (0.1% formic acid and 2% acetonitrile in water) at a concentration of 50 pM. The analysis of each peptide sample, with a loading amount of 50 pmol, was performed using a homemade C_18_ column via the auto-sampler integrated into the Easy-nLC 1100 system with a 10-minute gradient. Subsequently, the peptides were eluted and ionized under a voltage of 2.1 kV and detected using Q-Exactive mass spectrometry. The MS1 scan was detected by an Orbitrap analyzer with a resolution of 70,000 (m/z 200), covering a mass-to-charge (m/z) range from 450 to 1650. For the MS/MS scan, a resolution of 17,500 was employed. The automatic gain control of the full scan was set as 1×10^6^, the maximum injection time was set as 60 ms. The top five ions were selected for the fragmentation, utilizing a collision energy of 28%. The automatic gain control was set as 5×10^5^ and the dynamic exclusion time was set as 10s.

### Histone extraction and in-gel digestion

MCF7 cells were cultured in DMEM medium supplemented with 10% of FBS, 0.1% of penicillin and streptomycin stock. Cells were treated with 10 μM SAHA or JIB04 for 24 h and washed with cold PBS for 3 times before harvest. Histones were prepared as previously reported. Briefly, cells were lysed with extraction buffer (0.34 M sucrose, 10 mM KCl, 1.5 mM MgCl_2_, 10 mM HEPES, 5 mM sodium butyrate, 5 mM nicotinamide, 1x Protease inhibitor cocktail, supplemented with 0.5% NP-40). Pellets were collected after centrifugation and resuspended by 0.4N H_2_SO_4_. After incubation overnight, supernatant was collected and histones were precipitated by adding TCA to 20%. Histone sediment was washed by cold acetone (supplemented with 0.1% HCl) once and cold acetone twice. Samples were resolved by ddH_2_O and quantified by BCA assay. Samples were separated by SDS-PAGE and bands of core histones were excised. Trypsin was added and incubated at 37°C overnight for digestion. Tryptic histone peptides were collected and cleaned up by Zip-tip C18 for LC-MS/MS analysis.

### LC-MS/MS analysis for histone samples

Tryptic histone peptides were dissolved in buffer A (0.1% formic acid, H_2_O) and loaded onto C18 analytical column (ReproSil-Pur 120 C18-AQ, 1.9 μm particle size, 120 Å pore size, Dr. Maisch GmbH, Germany) via Vanquish Neo UHPLC system. Peptides were eluted by a linear gradient of increasing buffer B (0.1% formic acid, 90% ACN) and ionized for Orbitrap Ascend Tribrid Mass Spectrometer detection. Full scan was analyzed in Orbitrap under the resolution of 120K. MS/MS scan was performed in TopN mode. The top 10 precursors with charges +1 to +5 were selected by quadrupole. Selected precursors were fragmented by higher-energy collisional dissociation (HCD) with a stepped NCE of 20%, 25%, 30%. MS/MS scan was also analyzed in Orbitrap under the resolution of 15K. Dynamics exclusion was set as 10 s.

### Western blot analysis

Histone samples were separated by SDS-PAGE and transferred to the NC membrane. The member was blocked with 5% BSA and incubated with antibody overnight at 4°C. After washing with PBST (PBS supplemented with 0.1% Tween 20), the member was further incubated with second antibody. Then the member was washed with PBST again and detected by Tanon 5200 system.

### Data processing

For individual or mixed synthesized IMP datasets, raw MS files were converted to MGF/MS1/MS2 formats using pParse^45^. The MGF files were subsequently searched using Mascot software (version 2.6.0) against the human histone H4 sequence concatenated with E. coli protein sequences as background, which were downloaded from the UniProt database (updated on 08/18/2023). Acetyl (K), Oxidation (M), Propionyl-Methylation (K), Propionyl (K), and Propionyl (N-term) were specified as variable modifications. Mass tolerance was set to ± 10 ppm for precursor ions and ± 0.02 Da for the fragment ions. Trypsin was specified as the enzyme, allowing a maximum of 5 missed cleavages. The results were processed with custom in-house scripts to apply FDR filtering (threshold = 0.01) and convert to PTMdecoder-compatible input formats. Only MS/MS spectra with +2 charged precursor and >= 8 matched fragment ions were considered for single-peptidoform datasets. In PTMdecoder, the regularization coefficient lambda was set to 0.5. IMPs with relative abundance less than 10% of the maximum relative abundance in the same multiplexed spectrum were filtered out. Only methylation (+14.015650 Da) was permitted on lysine (K) residues in the C-terminal region of the peptide.

For the datasets of MCF7 cells treated with JIB-04/SAHA or DMSO, raw MS files were converted to MGF/MS1/MS2 formats using pParse. The MGF files were subsequently searched using Mascot software (version 2.6.0) against human histone sequences concatenated with E. coli protein sequences as background, which were downloaded from the UniProt database (updated on 08/18/2023). Acetyl (K), Dimethyl (K), Methyl (K), Trimethyl (K), and Oxidation (M) were specified as variable modifications. The results were converted into the input format for PTMdecoder using in-house codes. Peptide-spectrum matches (PSMs) were retained only if any N-terminal modification on the peptide was exclusively restricted to methylation, and were further filtered to ensure a maximum of two distinct modification types per peptide. Other parameters and processing steps were the same as those used for the synthesized-peptide datasets above. The normalization at both the peptidoform and site levels was performed manually using Excel.

### General model for deconvoluting multiplexed MS/MS spectra

In the MS/MS-level analysis (Fig. 1b), PTMdecoder takes the peptide identifications from traditional search engines, such as Mascot, as input. It first enumerates all possible IMPs according to the peptide sequence and PTM settings, then matches these IMPs to the multiplexed spectra. In the multiplexed spectra, only a subset of the fragment ions, called site-discriminating ions, are informative on the types and positions of modifications and are utilized in the model for deconvolution (Fig. 2c). Unlike some existing algorithms that rely solely on unique ions, PTMdecoder considers all site-discriminating ions, including shared ions across IMPs, which is essential for accurate IMP quantification in complex scenarios. Even when IMPs lack uniquely supporting fragment ions, shared fragment ions can nevertheless characterize the presence or absence of a set of IMPs. The combinatorial state of these fragment ions enhances the discrimination and quantification of individual IMPs. Since the peak intensities in multiplexed MS/MS spectra represent linear combinations of the peak intensities in individual IMPs, employing a non-negative linear model for deconvolution is natural and effective.

The general core model is established using site-discriminating ions as follows. Assuming there are 𝑁 site-discriminating fragment ions and 𝑚 potential IMPs, we build a linear model:

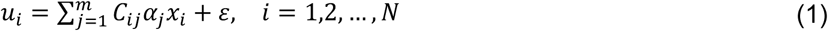

where 𝑢_𝑖_ is the observed intensity of the 𝑖th site-discriminating fragment ion (𝑠𝑑𝑓𝑖_𝑖_) in the spectrum, *m* is the total number of IMPs, 𝛼_𝑗_ is the abundance of the 𝑗th IMP (𝐼𝑀𝑃_𝑗_), 𝑥_𝑖_ is the fragmentation efficiency of 𝑠𝑑𝑓𝑖_𝑖_, and 𝐶_𝑖𝑗_ is an indicator function indicating whether 𝐼𝑀𝑃_𝑗_ can produce 𝑠𝑑𝑓𝑖_𝑖_:

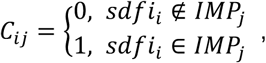

the error term ε is assumed to follow a zero-mean distribution.

In Equation (1), 𝐶_𝑖𝑗_ and 𝑢_𝑖_ are known, 𝑥_𝑖_ depends on the model assumption (unknown in PTMdecoder’s variable model and known in other two baseline models), and the objective is to solve 𝛼_𝑗_.

Since the abundance of an IMP cannot be negative, the fragmentation efficiencies should be in the range of 0 to 1. Based on these principles, the following constraints are used:

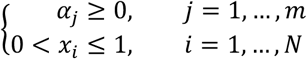

In PTMdecoder, the variable model and the two benchmark models all utilize this core model, differing primarily in their treatment of the fragmentation efficiency 𝑥_𝑖_.

### Variable model of fragmentation efficiency

In the variable model of fragmentation efficiency, fragment ions sharing the same terminal type, number of residues, and charge state are grouped into the same class. As previous works and our experimental results have demonstrated (Fig. 2, Supplementary Fig. 3-4), the fragmentation efficiencies of ions within the same class are similar among IMPs regardless of the presence or absence of modifications, whereas significant differences may occur across different classes^22,28^. Therefore, a single variable can be assigned to each ion class, rather than modeling individual ions separately.

Assuming that the fragment ions can be divided into 𝑞 classes, the fragmentation efficiency 𝑥_𝑖_ (𝑖 = 1, …, 𝑁) can be rewritten as 𝑡_𝑘𝑖_ (𝑘_𝑖_ = 1, …, 𝑞). The subscript 𝑖 added in 𝑘_𝑖_ indicates that the value of 𝑘 is determined by the classification of 𝑥_𝑖_. This reduces the number of unknown variables in Equation (1) from 𝑚 + 𝑁 to 𝑚 + 𝑞. By moving the fragmentation efficiency variables to the left-hand side, Equation (1) can be expressed in linear form:

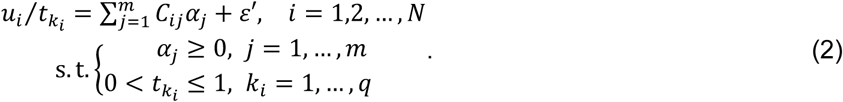

Furthermore, it can be represented in matrix notation by defining:

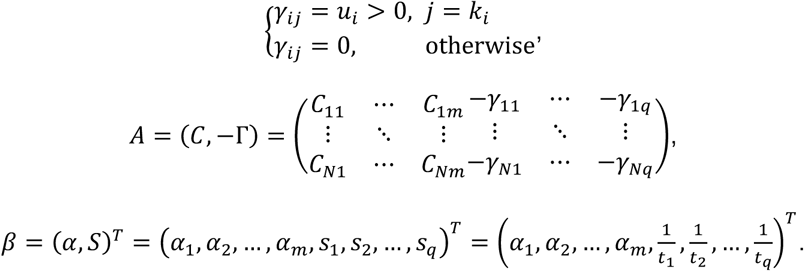

Equation (2) can be rewritten by moving the term 𝑢_𝑖_⁄𝑡_𝑘𝑖_ to the right-hand side:

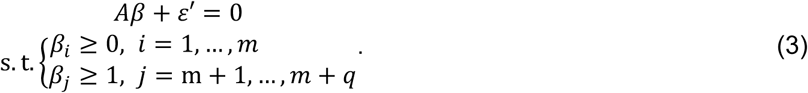

Given the negligible magnitude of systematic errors 𝜀′ relative to peptidoform-derived signal intensities, we determined the refined estimation of 𝛽 by minimizing the 𝐿_2_-norm of 𝐴𝛽:

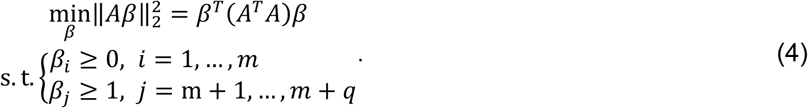

While Equation (4) provides a baseline estimate of IMP abundances by minimizing spectral fitting errors, its solutions tend to assign non-zero contributions to some implausible IMPs. To align the solutions with the empirical principle that only a subset of IMPs co-elutes in a real mixture, a 𝐿_1_regularization penalty is added to the model inspired by Lasso:

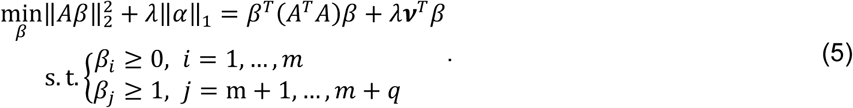

where 𝜆 > 0 is a penalty parameter controlling sparsity strength, 𝝂 = (𝟏_1×m_, 𝟎_1×𝑞_)^𝑇^ = (1, …,1,0,0, …,0)^𝑇^ is a binary mask vector that selectively applies the 𝐿_1_ penalty only to peptidoform abundances (𝛼_𝑖_, first 𝑚 components of 𝛽). This mechanism discriminates the composing peptidoforms in the mixture by driving the abundance of irrelevant peptidoform to exact zero. Notably, 𝐿_1_ regularization is also adopted in the two baseline models discussed below.

### Constant and equal models of fragmentation efficiency

The constant model of fragmentation efficiency also assumes that fragmentation ions can be grouped into classes according to the terminal type, number of residues, and charge state of fragment ions. However, unlike the variable model, the fragmentation efficiencies are pre-estimated before solving the model. Let 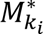 be the aggregate intensity of fragment ions in class 𝑘_𝑖_ in the spectrum 𝑀 be the total abundance of all IMPs. Then, the fragmentation efficiency of the class 𝑘_𝑖_ ions is 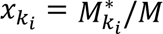. When the relative abundance is desired, 𝑀 can be ignored. Therefore, the only unknown variables in Equation (1) are the relative abundances of IMPs, i.e., 𝛼_𝑗_ and Equation (1) can be written as

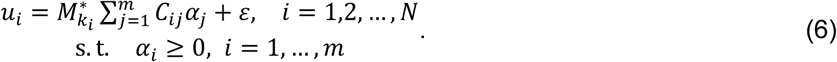

The equal model of fragmentation efficiency assumes that the fragmentation efficiencies of all ions are equal^28^. Therefore, Equation (1) can be rewritten as

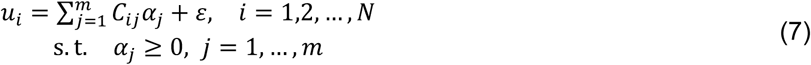

### Deconvolution of XIC peaks of co-eluted IMPs

Due to the dynamic exclusion in DDA, not all MS1 spectra are accompanied by an MS2 spectrum representing the target IMPs. Therefore, estimating the relative abundance of each IMP at each retention time is necessary for achieving precise quantification. Unlike previous methods that determine the IMP proportions at each retention time using linear interpolation^31^, PTMdecoder employs a non-parametric approach to estimate the relative abundance of each IMP at the retention times corresponding to acquired MS1 spectra (Supplementary Note 1). This approach accounts for the local characteristic of the quantification trend observed in the MS/MS spectra (Fig. 1d). The total XIC peak of the co-eluted IMPs is deconvoluted to several reconstructed XIC peaks, each corresponding to an individual IMP. PTMdecoder finally quantifies the IMPs by integrating the area under each reconstructed XIC curve.

## Acknowledgments

This work was supported by the National Key R&D Program of China (Grants 2022YFA1004801 and 2022YFA1304603), the National Natural Science Foundation of China (Grants 32070668, 22225702, T2488301, and 32171434). We thank Mingzhou Deng for his insights related to the manuscript.

## Supplementary Figures

**Supplementary Figure 1.**
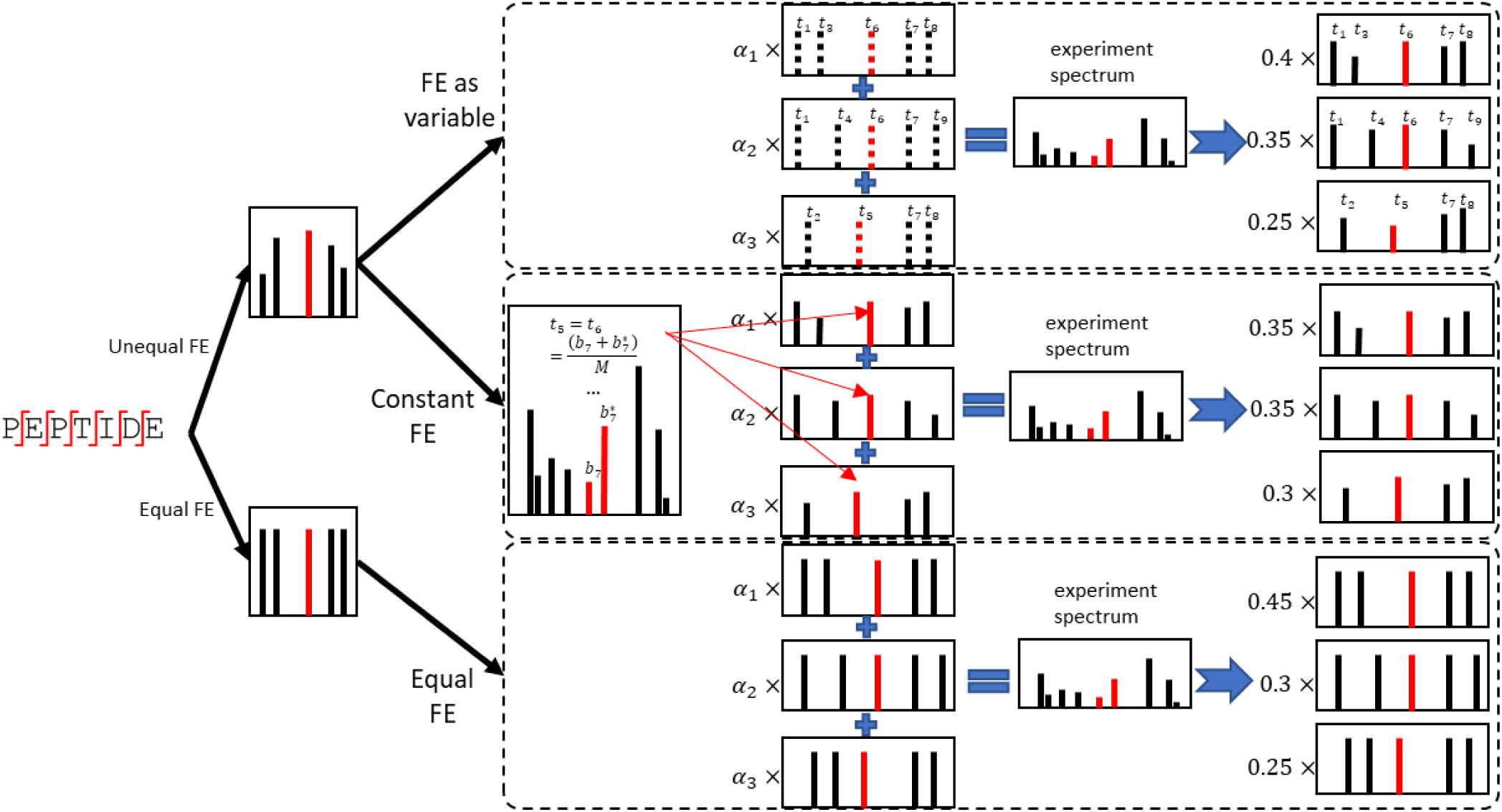
Schematic diagram of three models based on different utilization strategies of fragmentation efficiency (FE).

**Supplementary Figure 2a.**
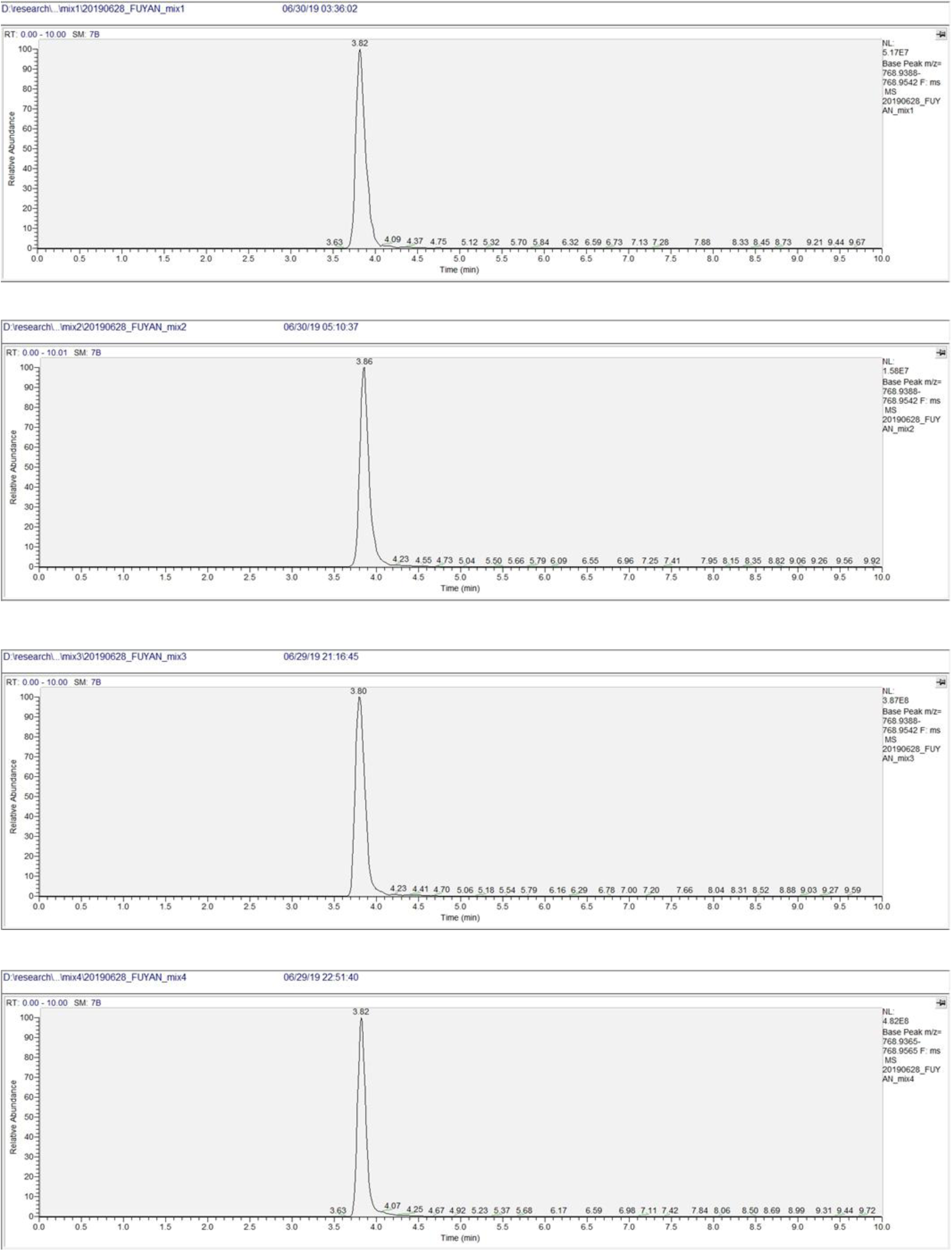
Most mixed samples produced single XIC peaks across the entire chromatographic gradient.

**Supplementary Figure 2b.**
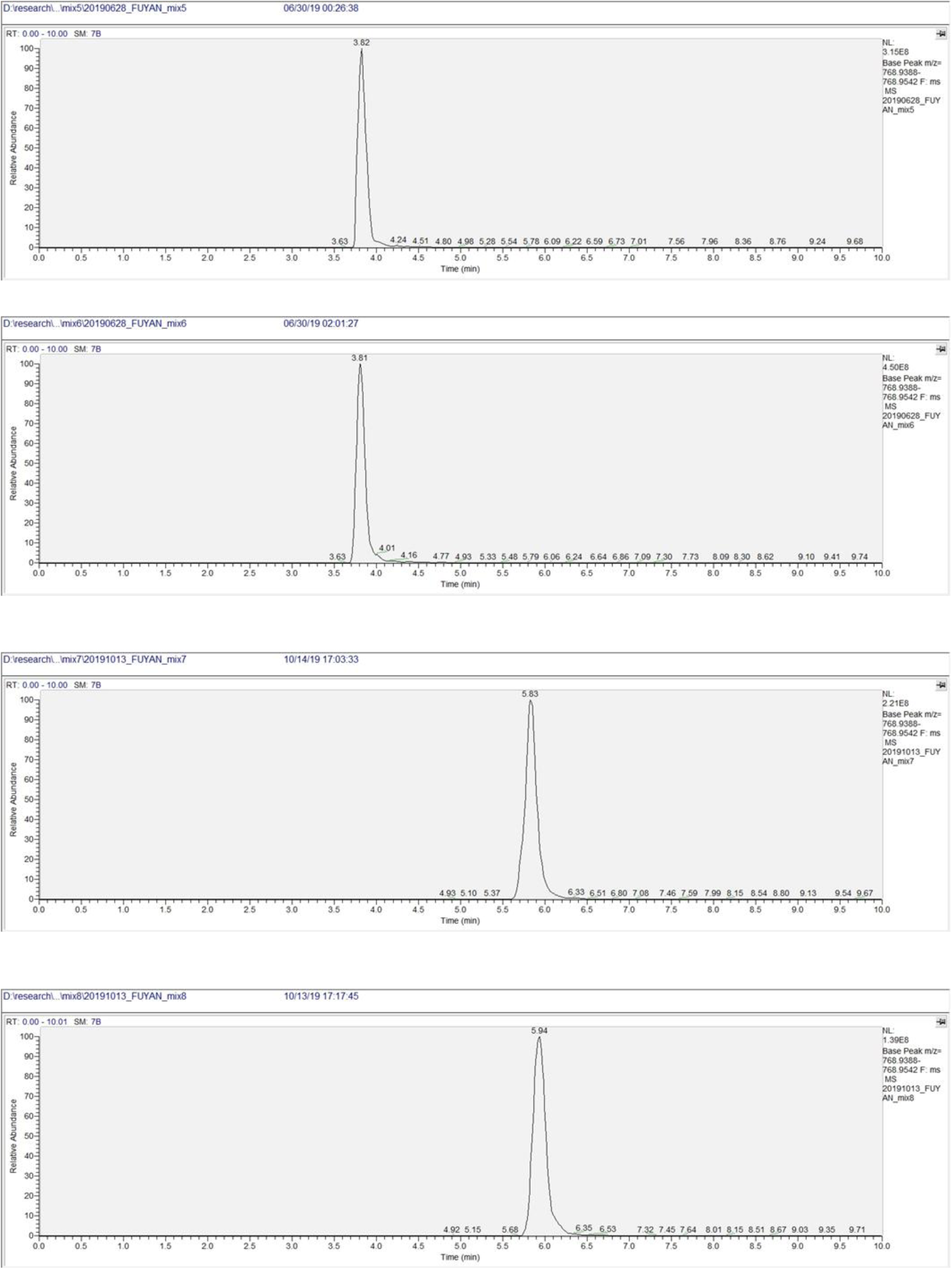
Most mixed samples produced single XIC peaks across the entire chromatographic gradient.

**Supplementary Figure 2c.**
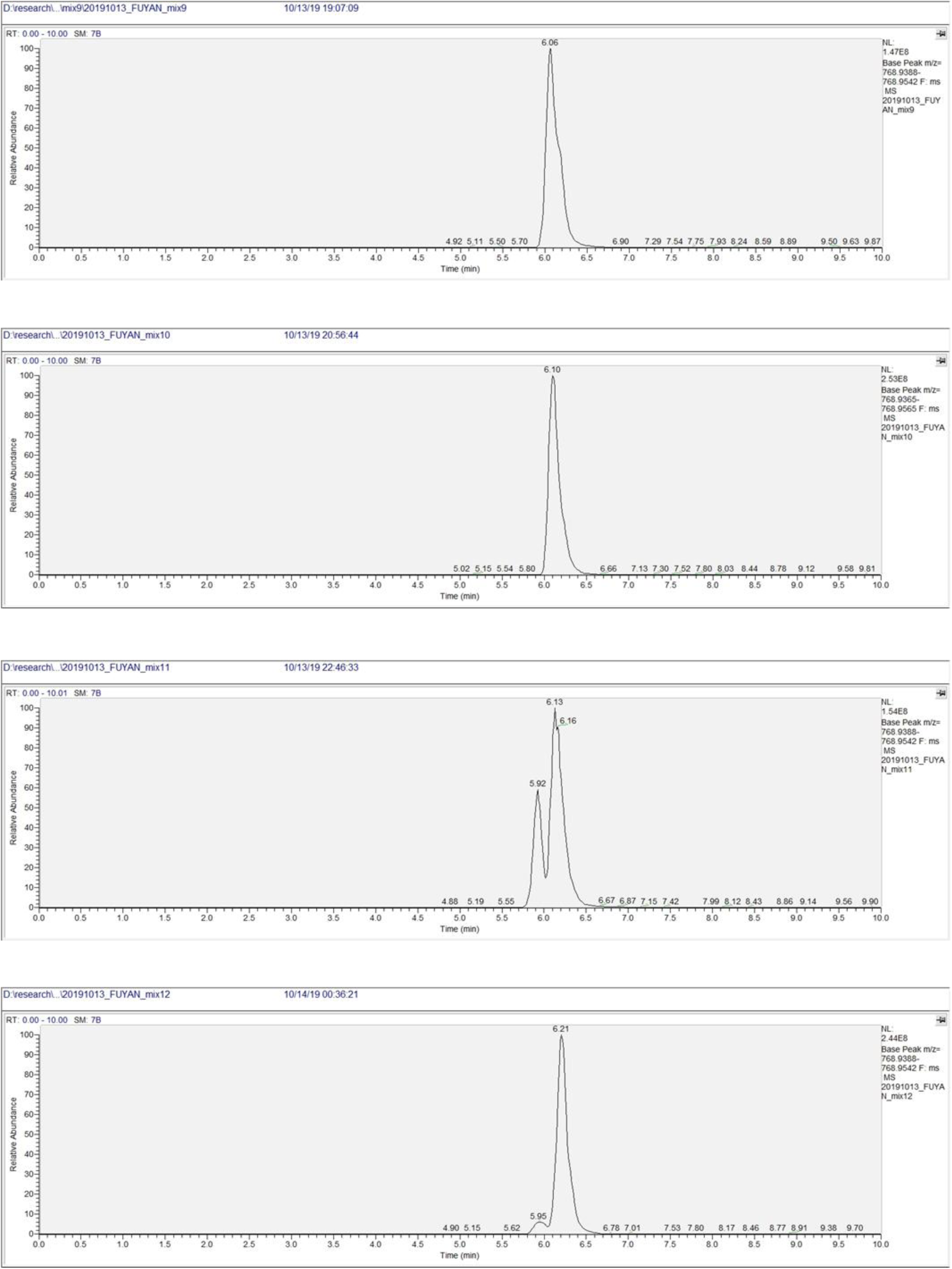
Most mixed samples produced single XIC peaks across the entire chromatographic gradient.

**Supplementary Figure 2d.**
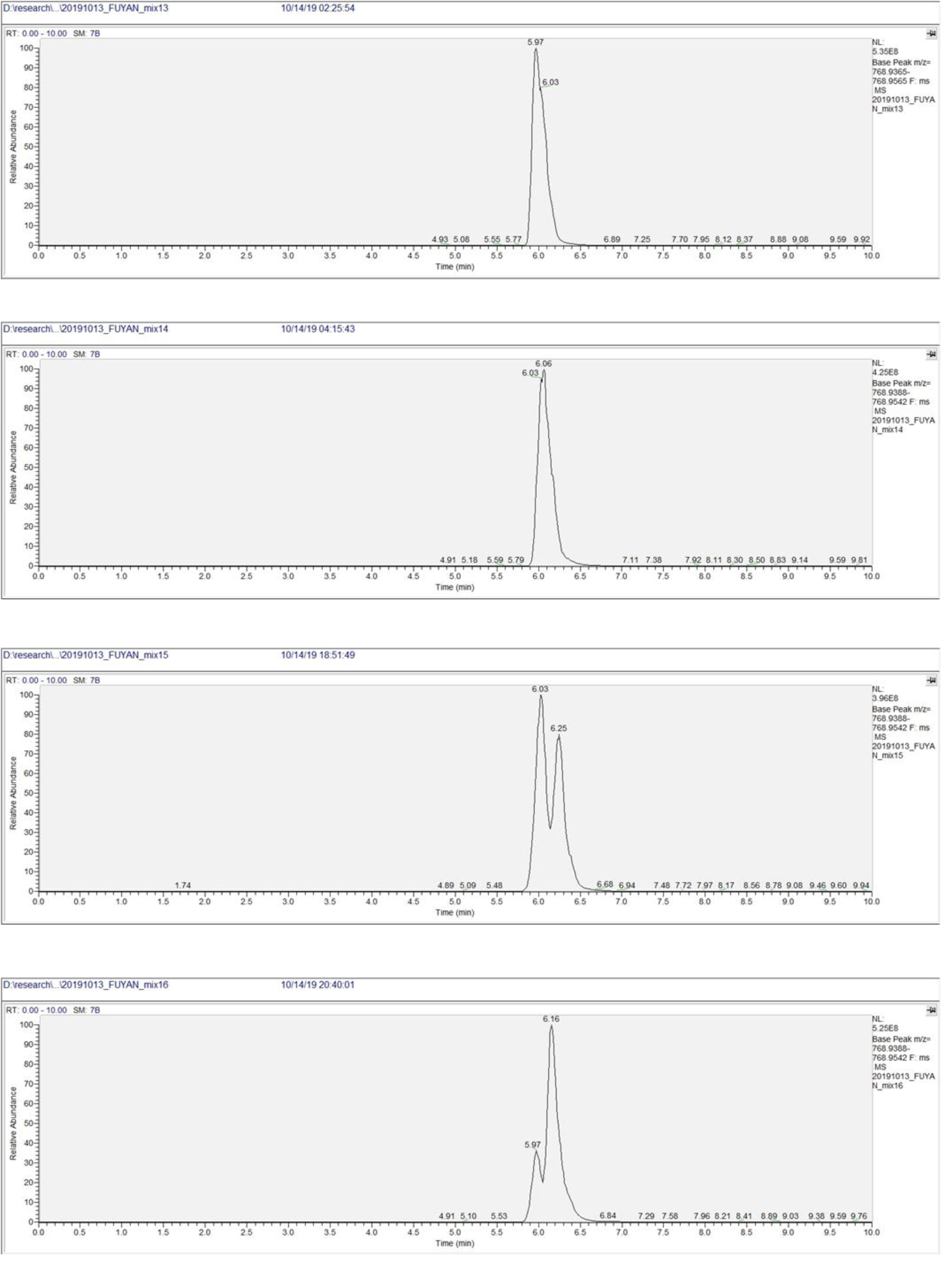
Most mixed samples produced single XIC peaks across the entire chromatographic gradient.

**Supplementary Figure 2e.**
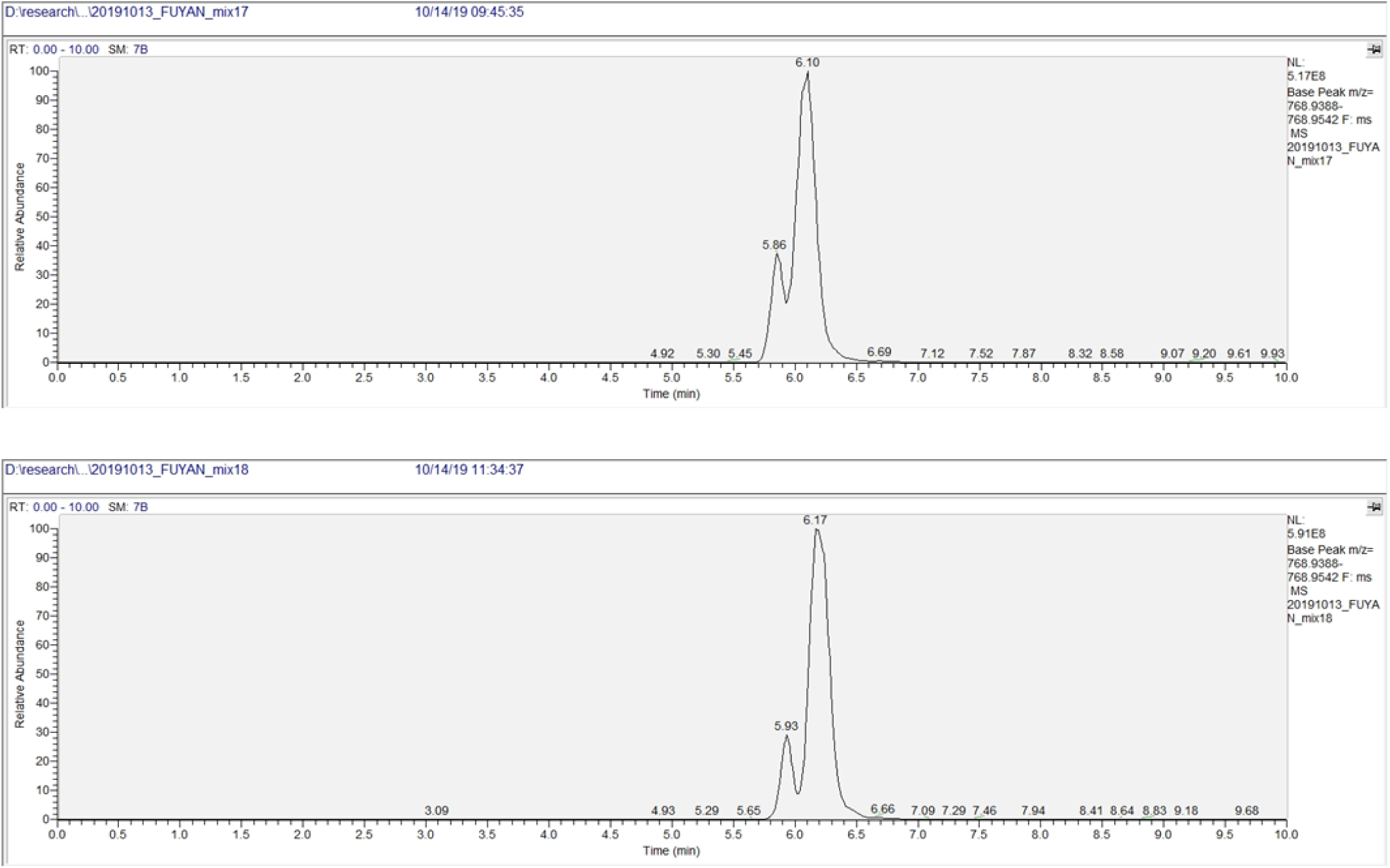
Most mixed samples produced single XIC peaks across the entire chromatographic gradient.

**Supplementary Figure 3.**
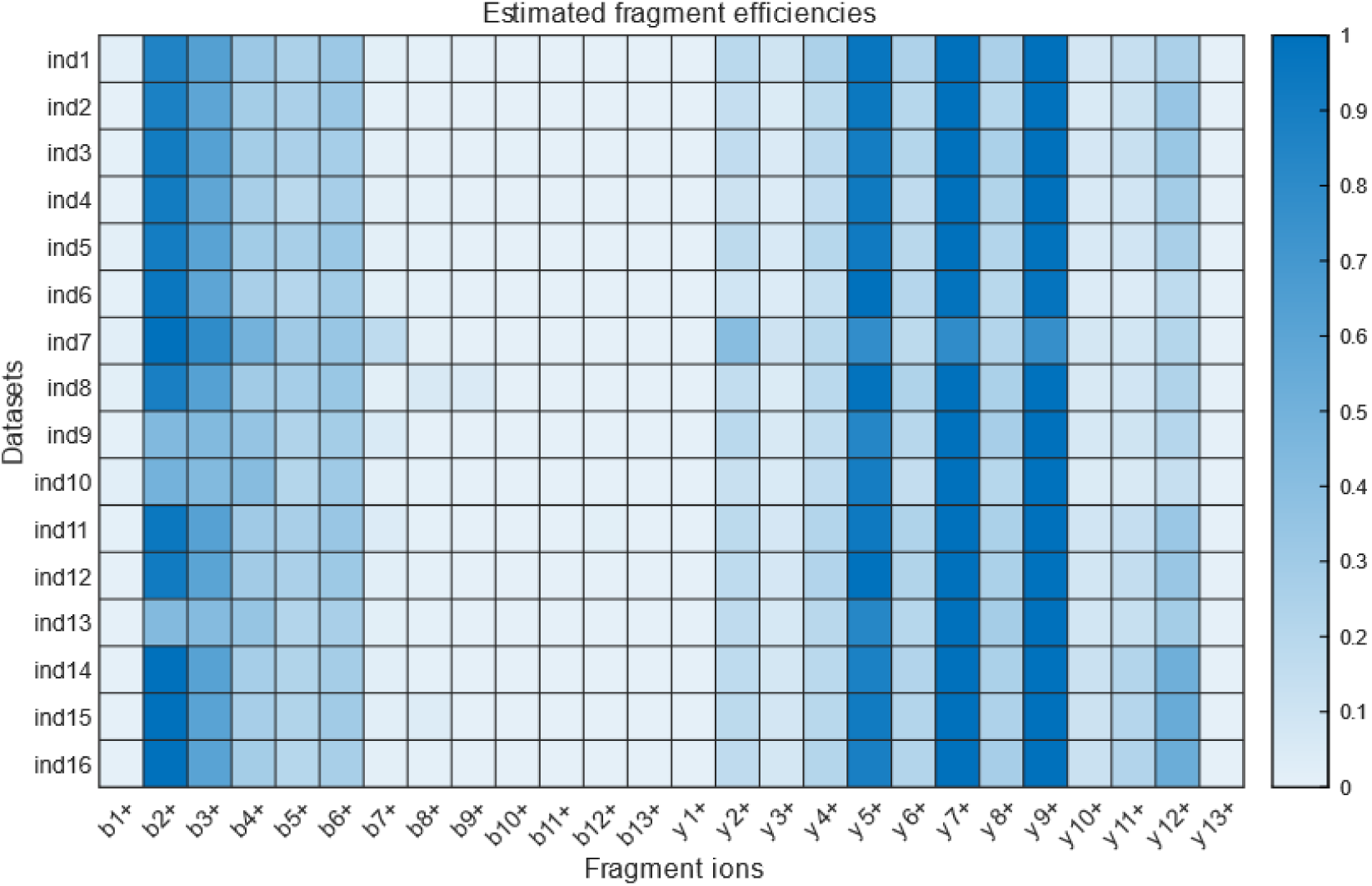
Heatmap of average estimated fragmentation efficiency for ion classes (x-axis) across 16 single-peptidoform datasets (y-axis).

**Supplementary Figure 4.**
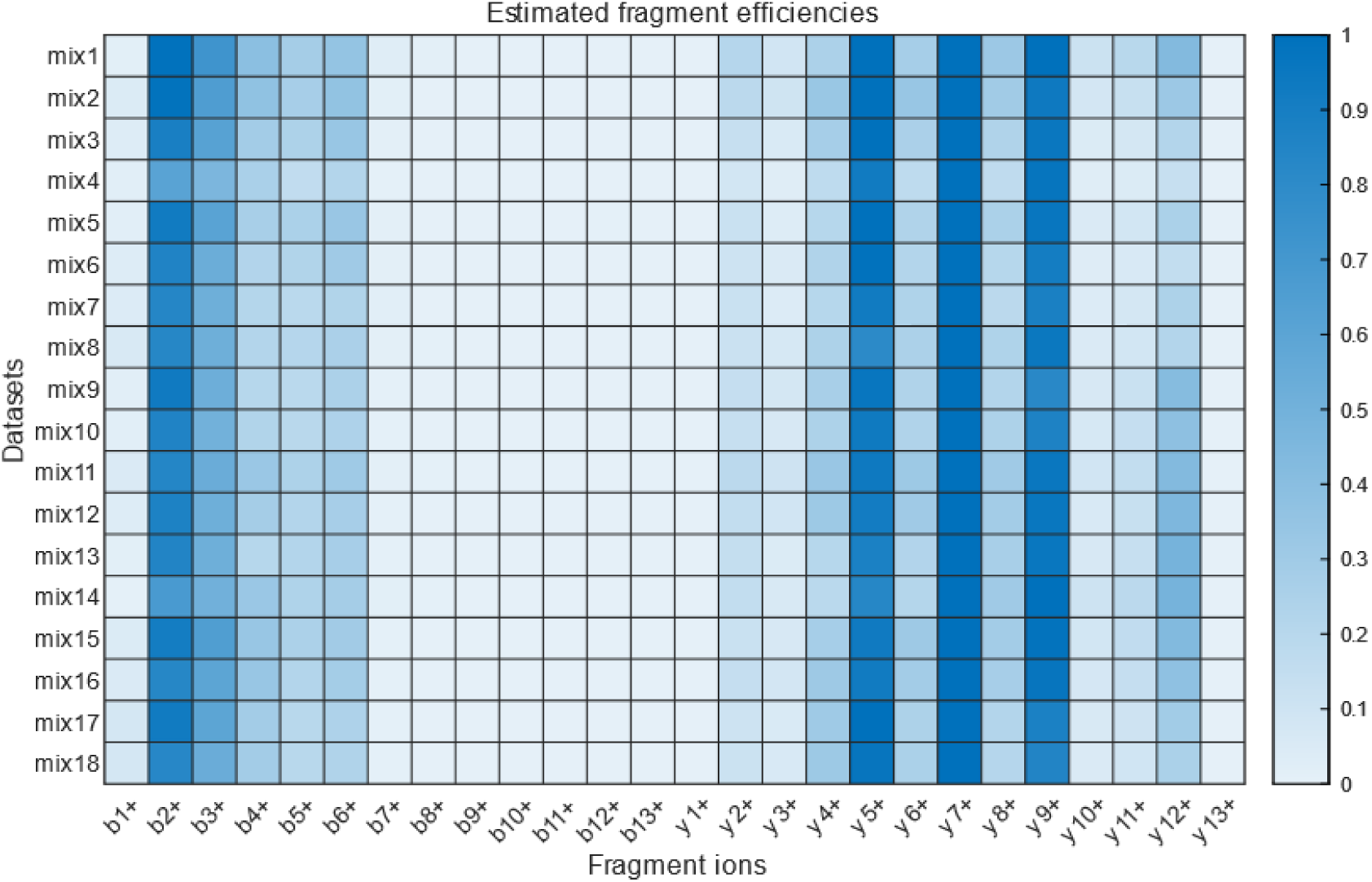
Heatmap of average estimated fragmentation efficiency for ion classes (x-axis) across 18 mixed-IMP datasets (y-axis).

**Supplementary Figure 5.**
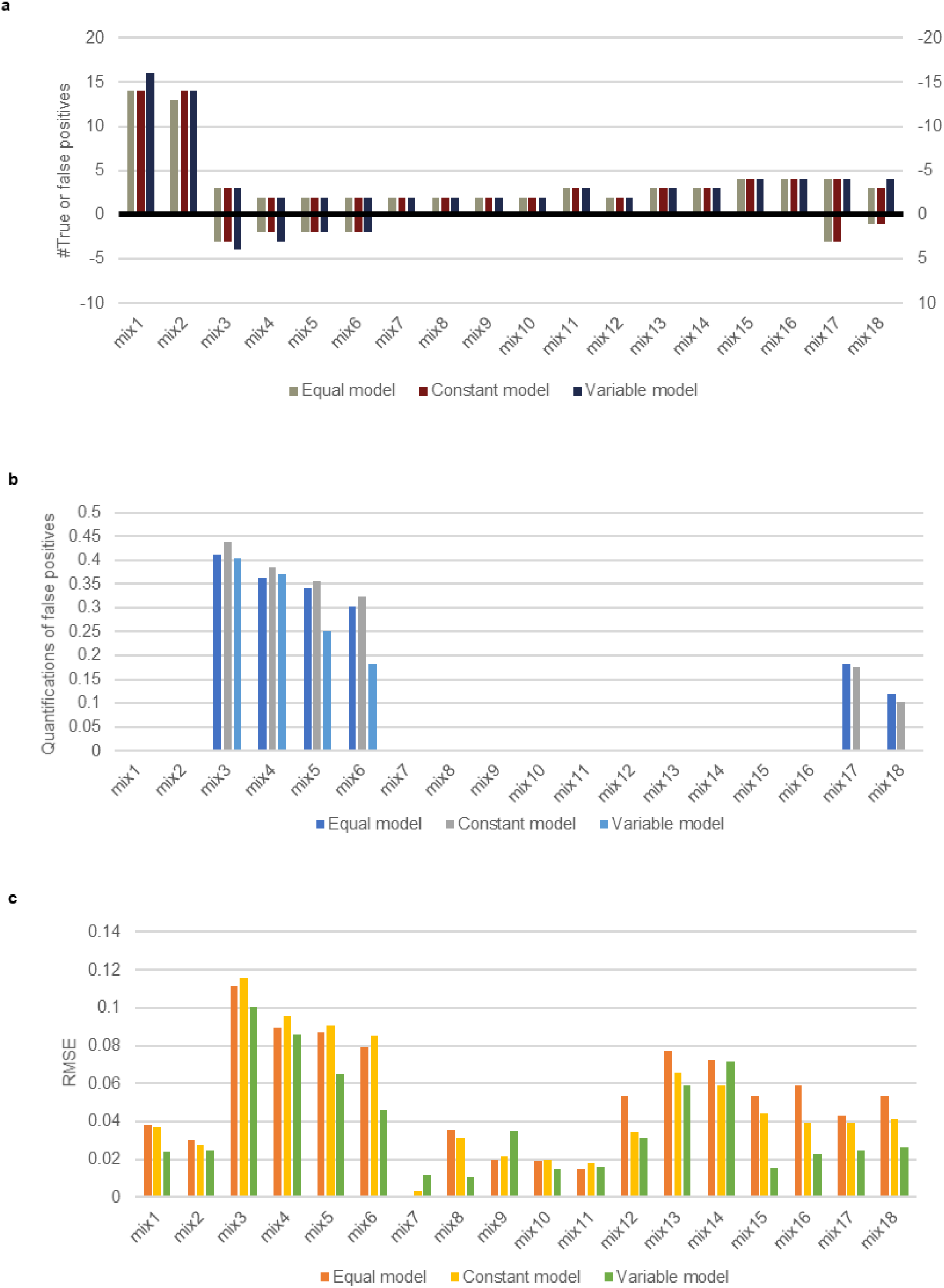
Performance comparison of three models for mixed-IMP samples. (a) Numbers of true positive and false positive. (b) Relative abundance of false positives. (c) RMSEs of quantification.

**Supplementary Figure 6a.**
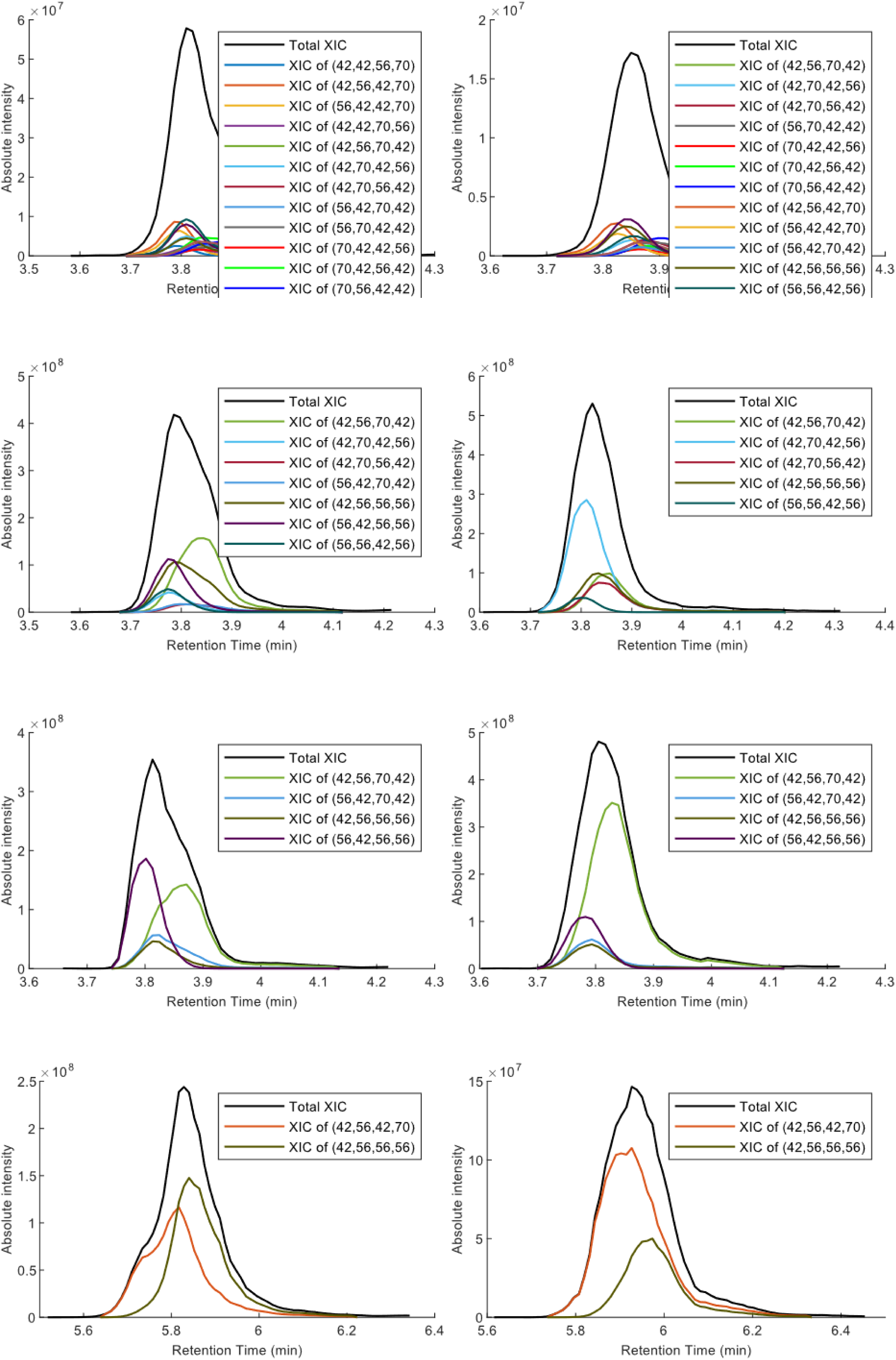
Deconvolution of total XIC in 18 mixed samples. (left) XICs of IMPs deconvoluted from the equally-mixed IMP datasets. (right) XICs of IMPs deconvoluted from the unequally-mixed IMPs datasets with same peptidoform components.

**Supplementary Figure 6b.**
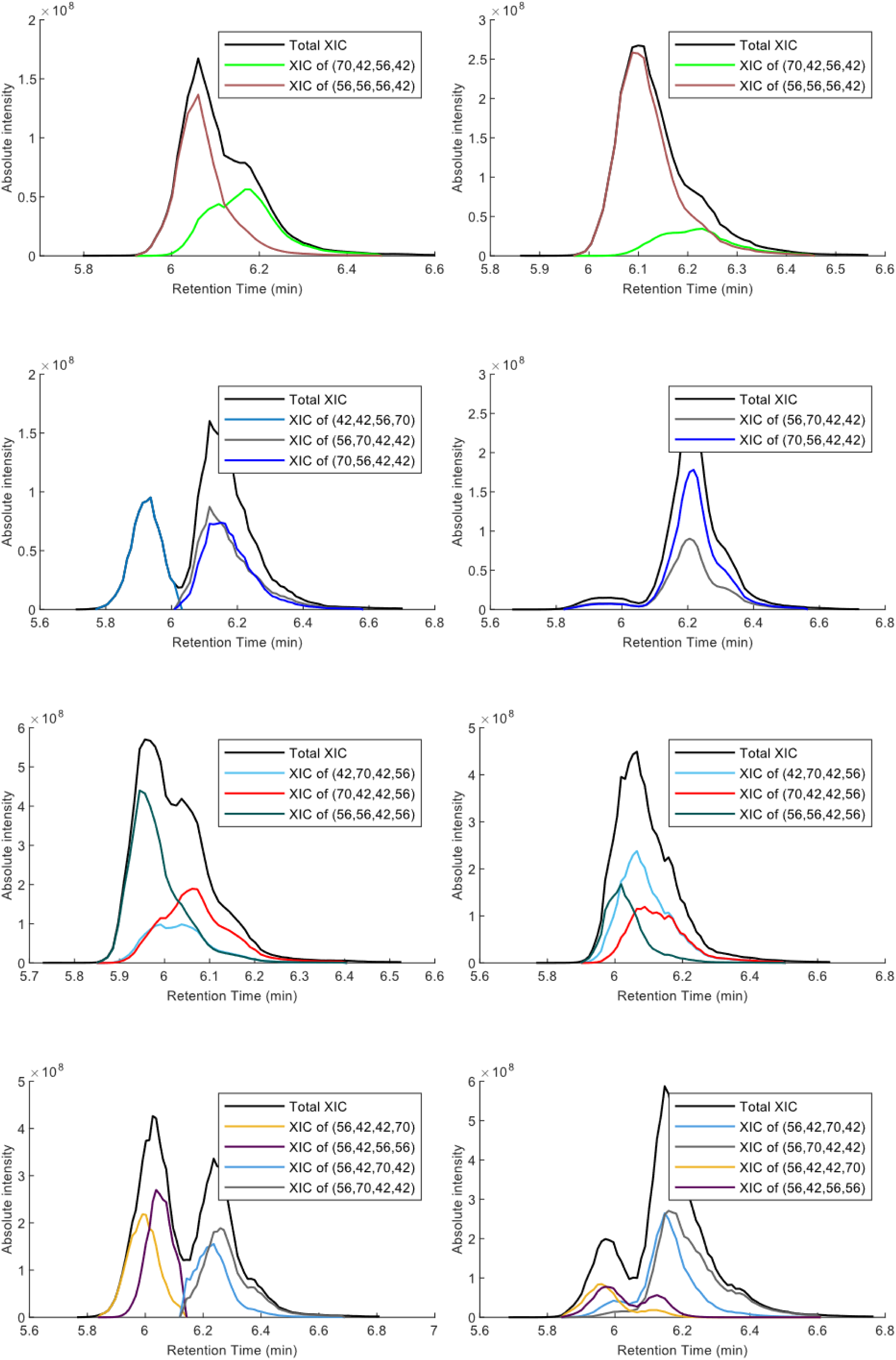
Deconvolution of total XIC in 18 mixed samples. (left) XICs of IMPs deconvoluted from the equally-mixed IMP datasets. (right) XICs of IMPs deconvoluted from the unequally-mixed IMPs datasets with same peptidoform components.

**Supplementary Figure 6c.**
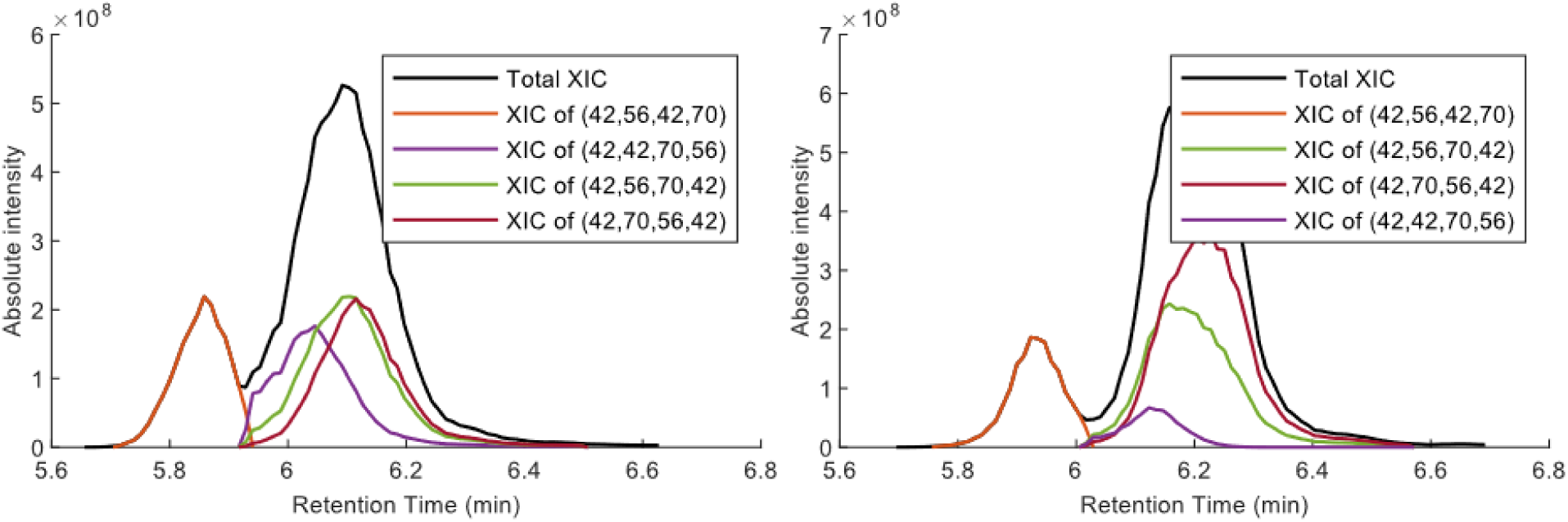
Deconvolution of total XIC in 18 mixed samples. (left) XICs of IMPs deconvoluted from the equally-mixed IMP datasets. (right) XICs of IMPs deconvoluted from the unequally-mixed IMPs datasets with same peptidoform components.

**Supplementary Figure 7a.**
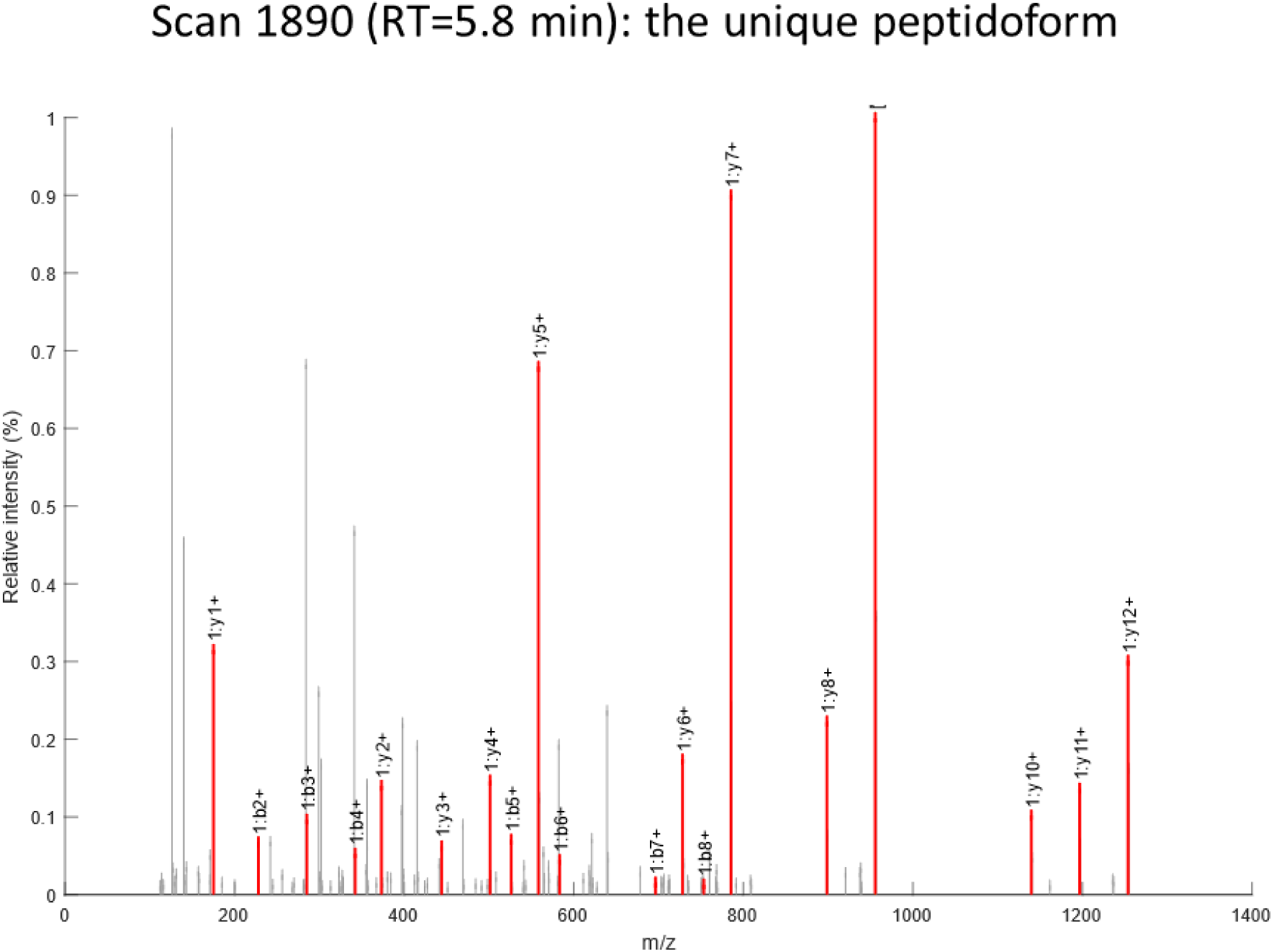
MS/MS spectra of co-eluting isobaric modification peptidoforms (IMPs) at four retention times in Fig. 3f. Top-scoring IMPs and all PTMdecoder-discriminated IMPs were labeled separately. Peaks are color-annotated: red (fragment ions unique to single peptidoforms), blue (shared fragment ions), green (precursor ions), gray (unmatched).

**Supplementary Figure 7b.**
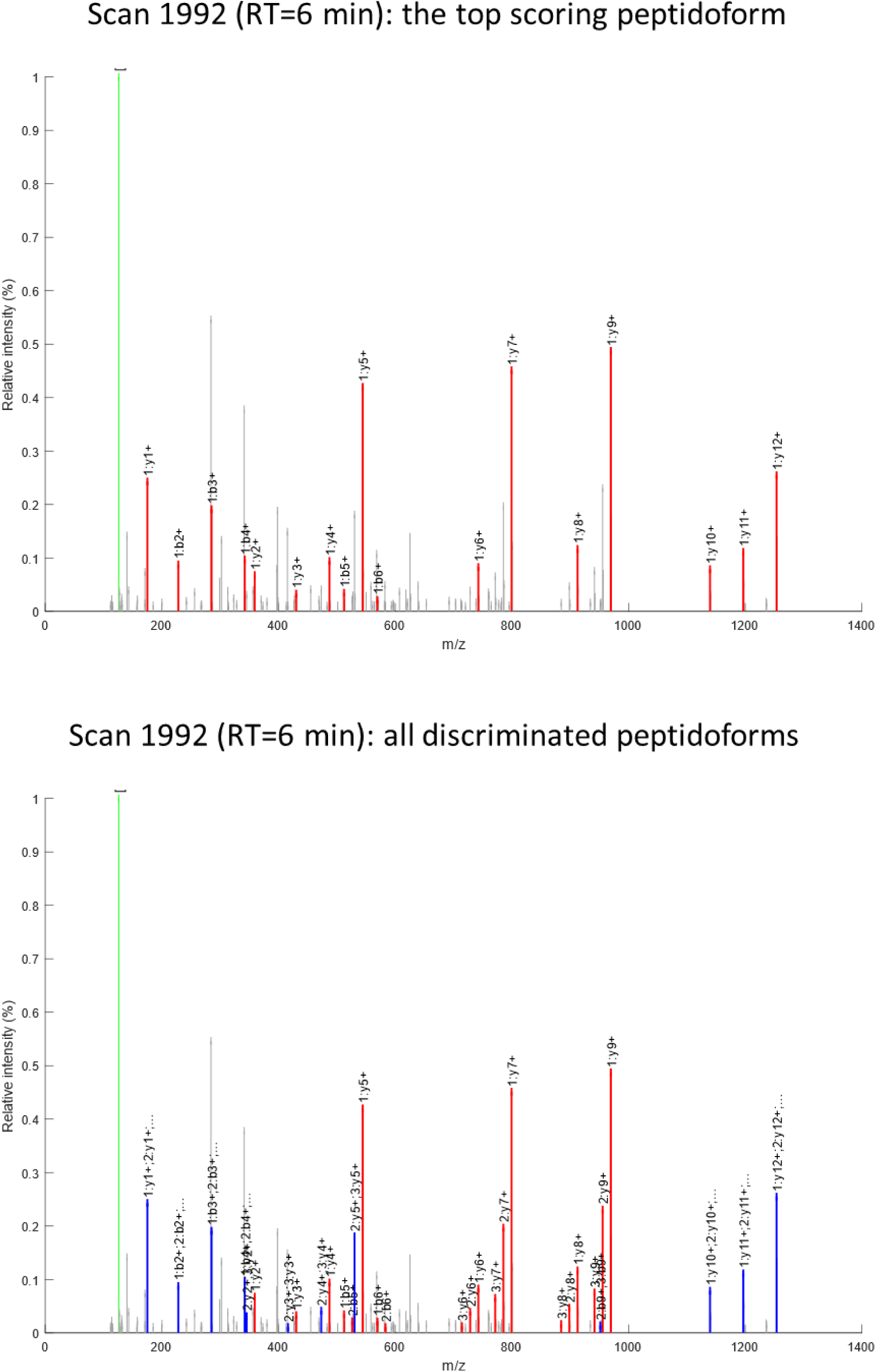
MS/MS spectra of co-eluting isobaric modification peptidoforms (IMPs) at four retention times in Fig. 3f.

**Supplementary Figure 7c.**
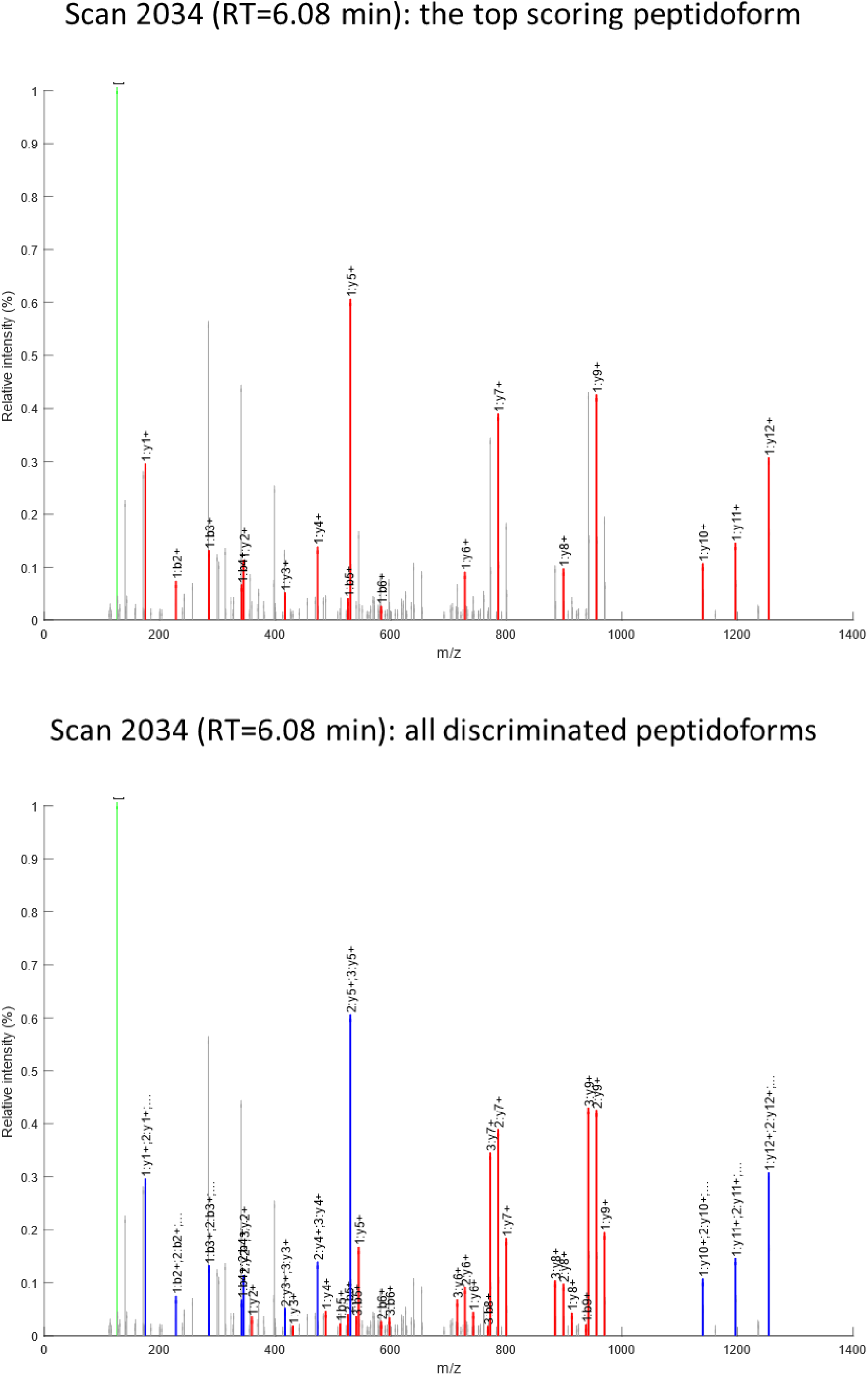
MS/MS spectra of co-eluting isobaric modification peptidoforms (IMPs) at four retention times in Fig. 3f.

**Supplementary Figure 7d.**
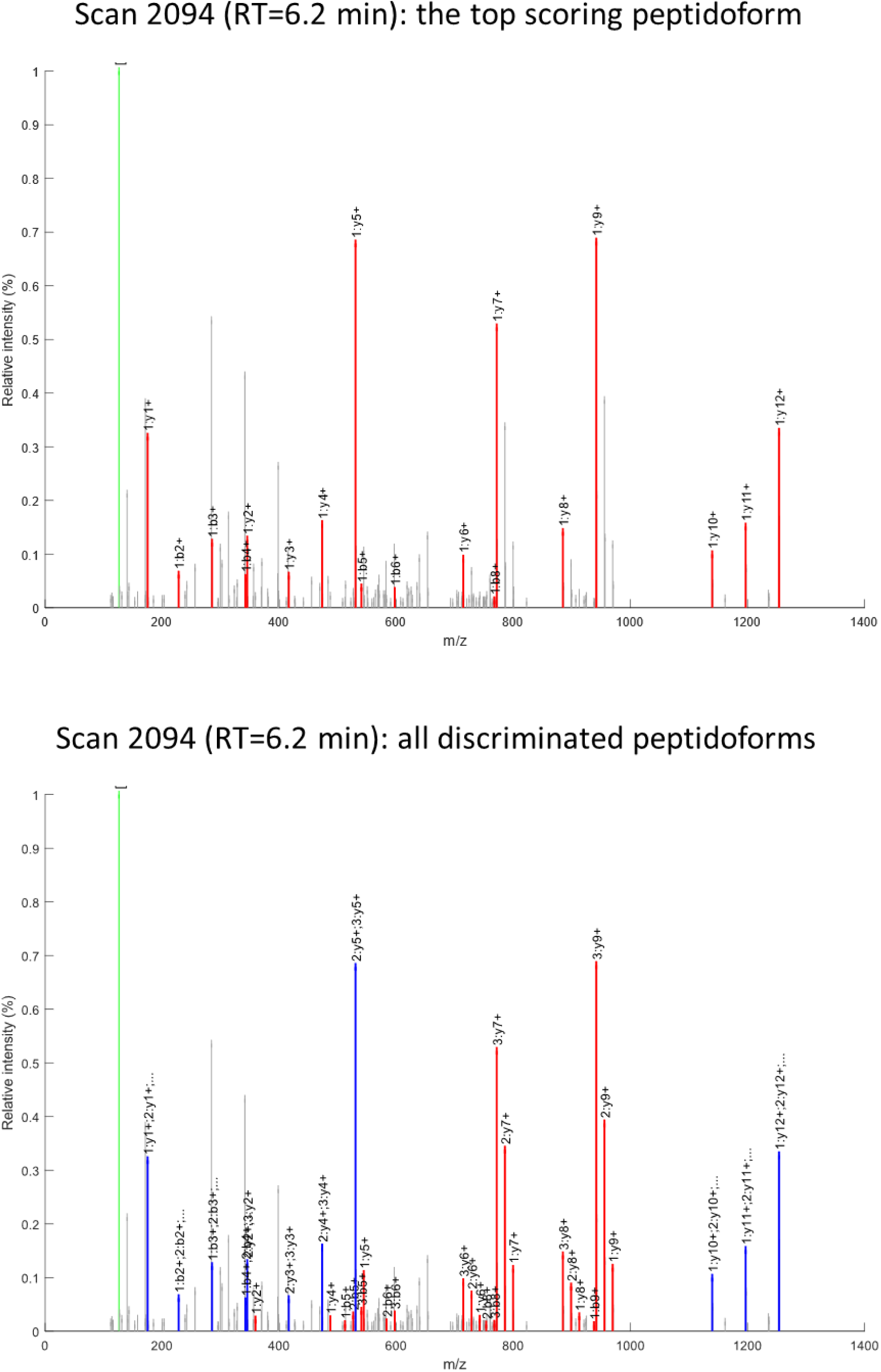
MS/MS spectra of co-eluting isobaric modification peptidoforms (IMPs) at four retention times in Fig. 3f.

**Supplementary Figure 8a.**
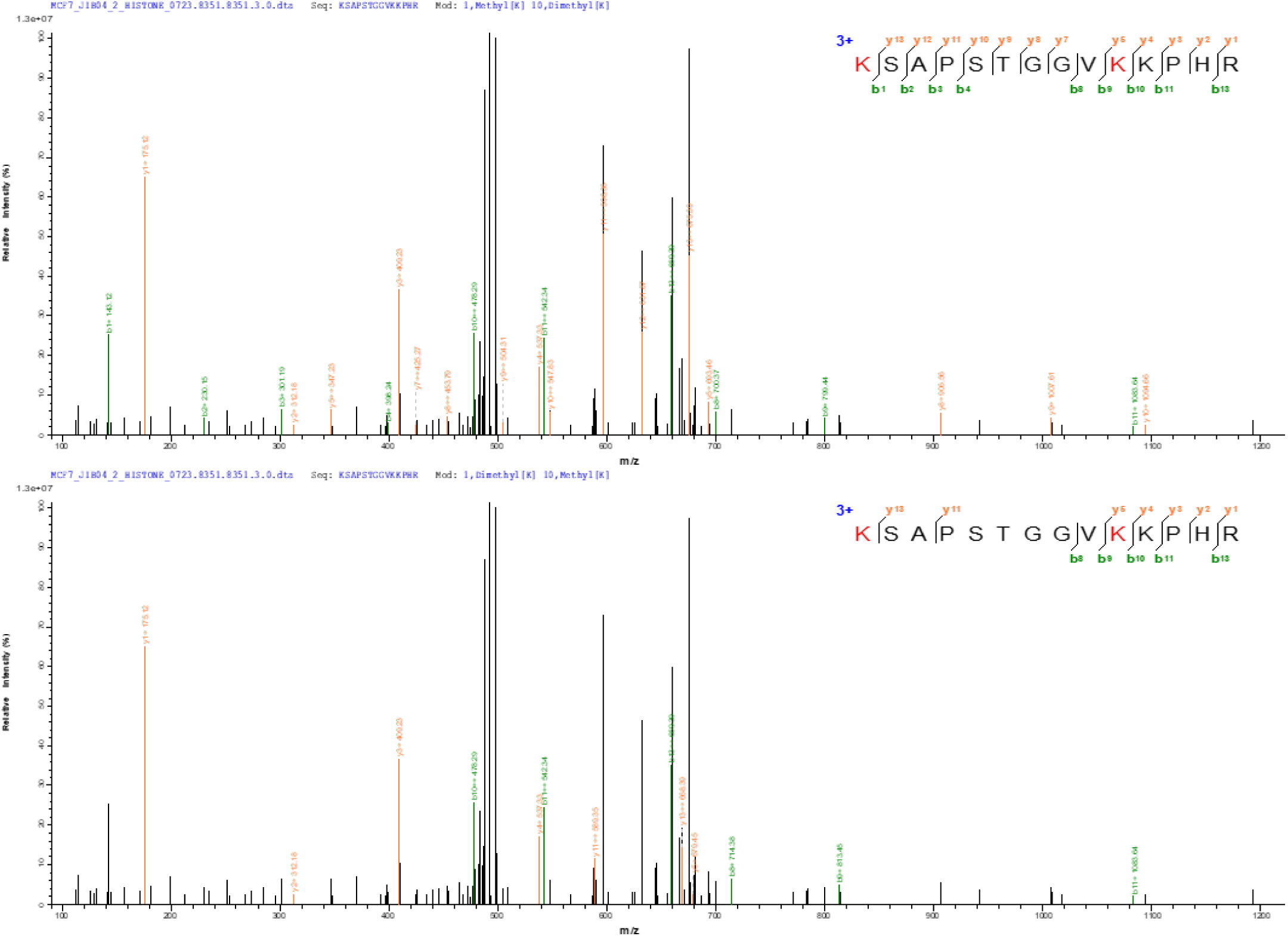
MS/MS spectral evidence supporting deconvolution in Fig. 4h. Annotated spectra spanning full XIC peak retention times, with peptidoform identifications confirming co-elution patterns.

**Supplementary Figure 8b.**
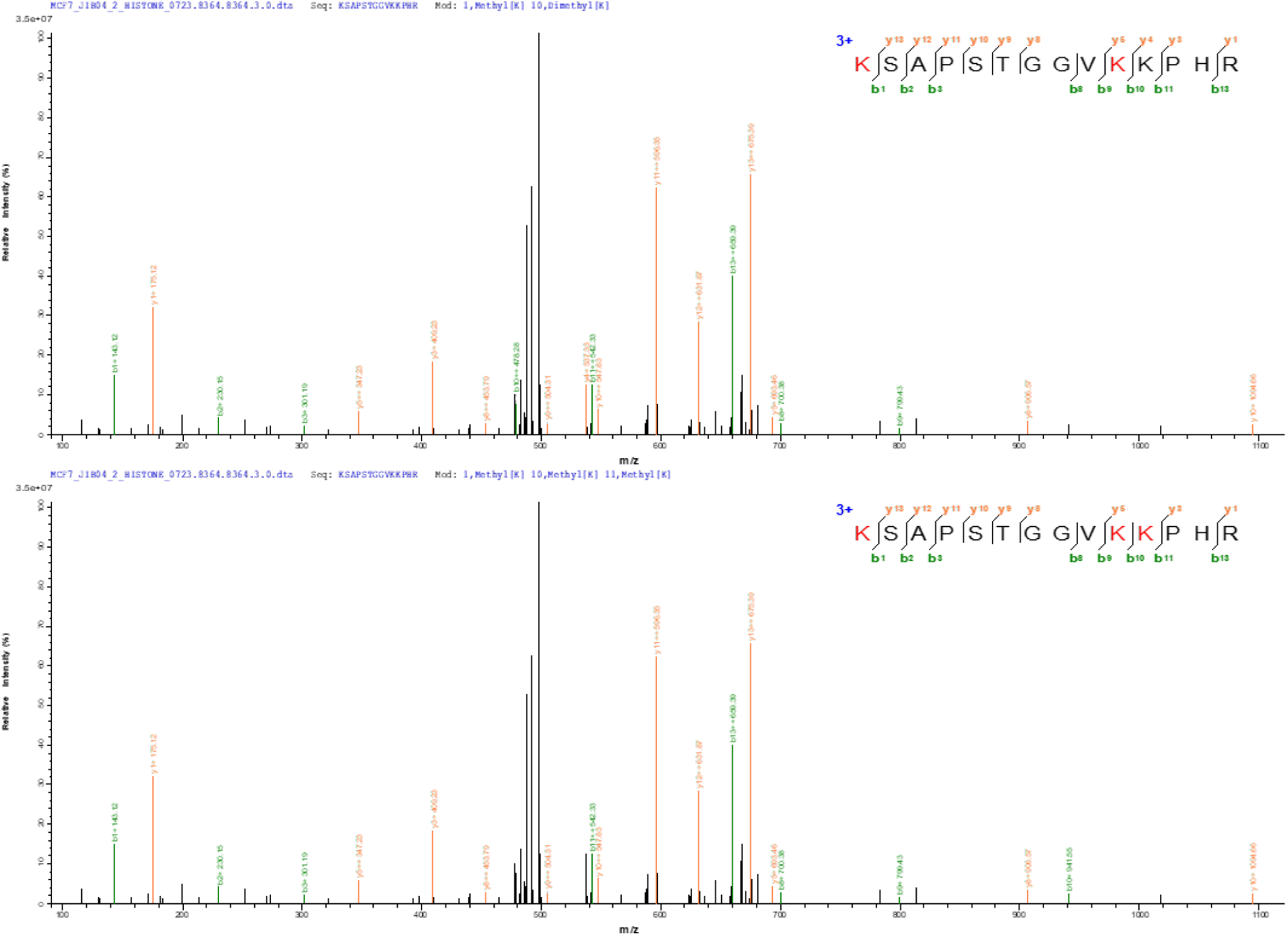
MS/MS spectral evidence supporting deconvolution in Fig. 4h.

**Supplementary Figure 8c.**
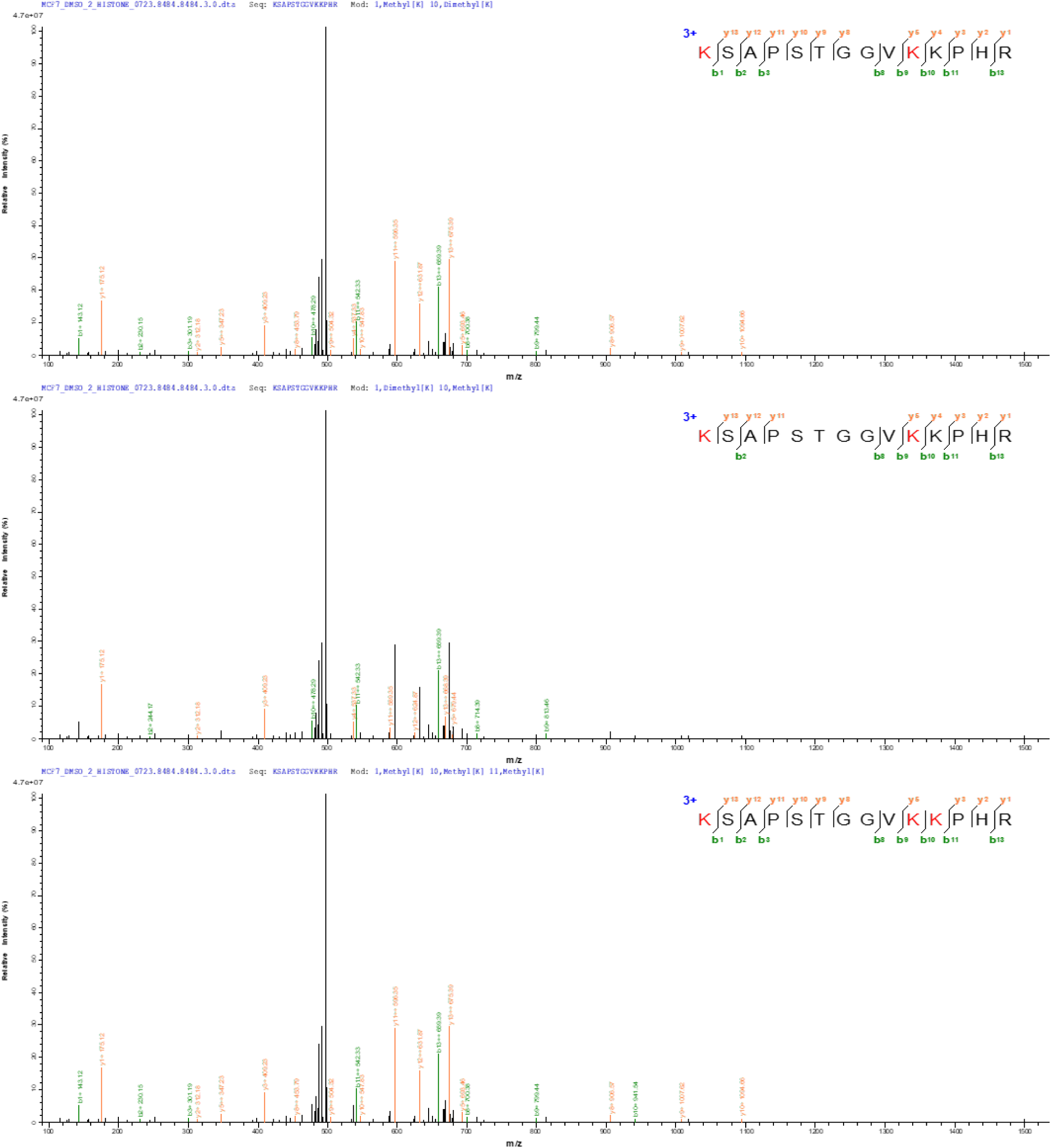
MS/MS spectral evidence supporting deconvolution in Fig. 4i.

**Supplementary Figure 9a.**
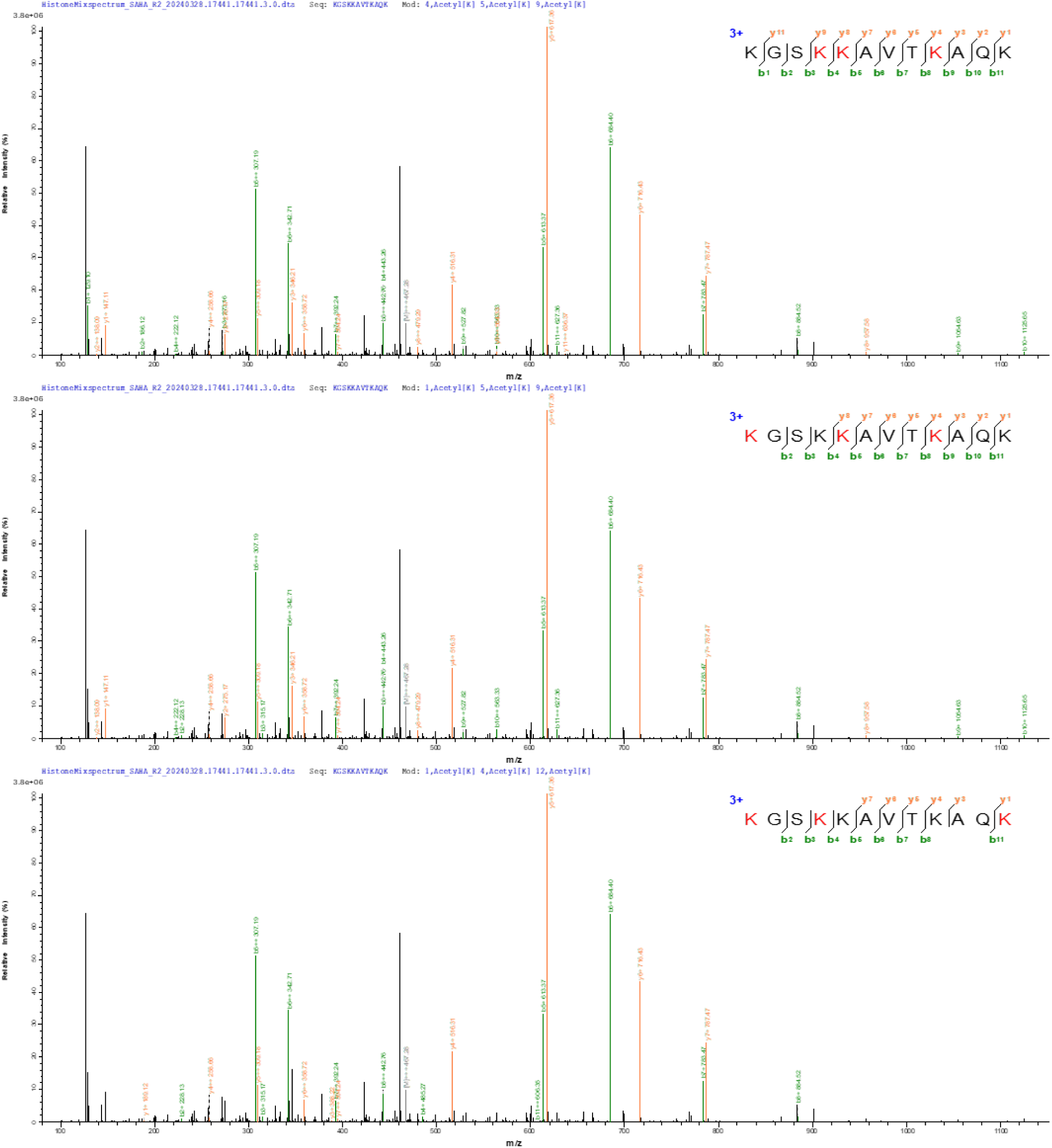
MS/MS spectral evidence supporting deconvolution in Fig. 5h. Annotated spectra spanning full XIC peak retention times, with peptidoform identifications confirming co-elution patterns.

**Supplementary Figure 9b.**
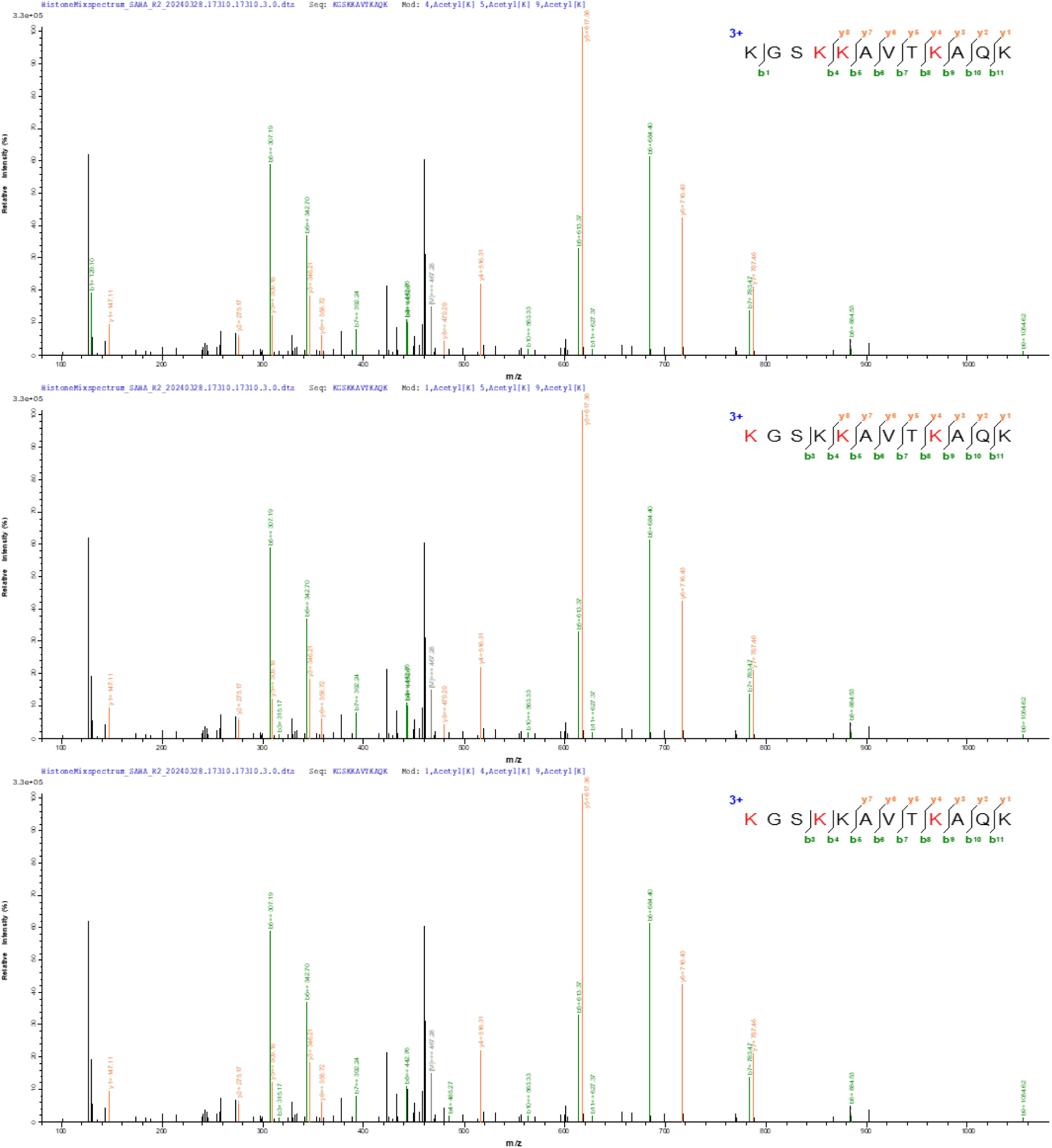
MS/MS spectral evidence supporting deconvolution in Fig. 5h.

**Supplementary Figure 9c.**
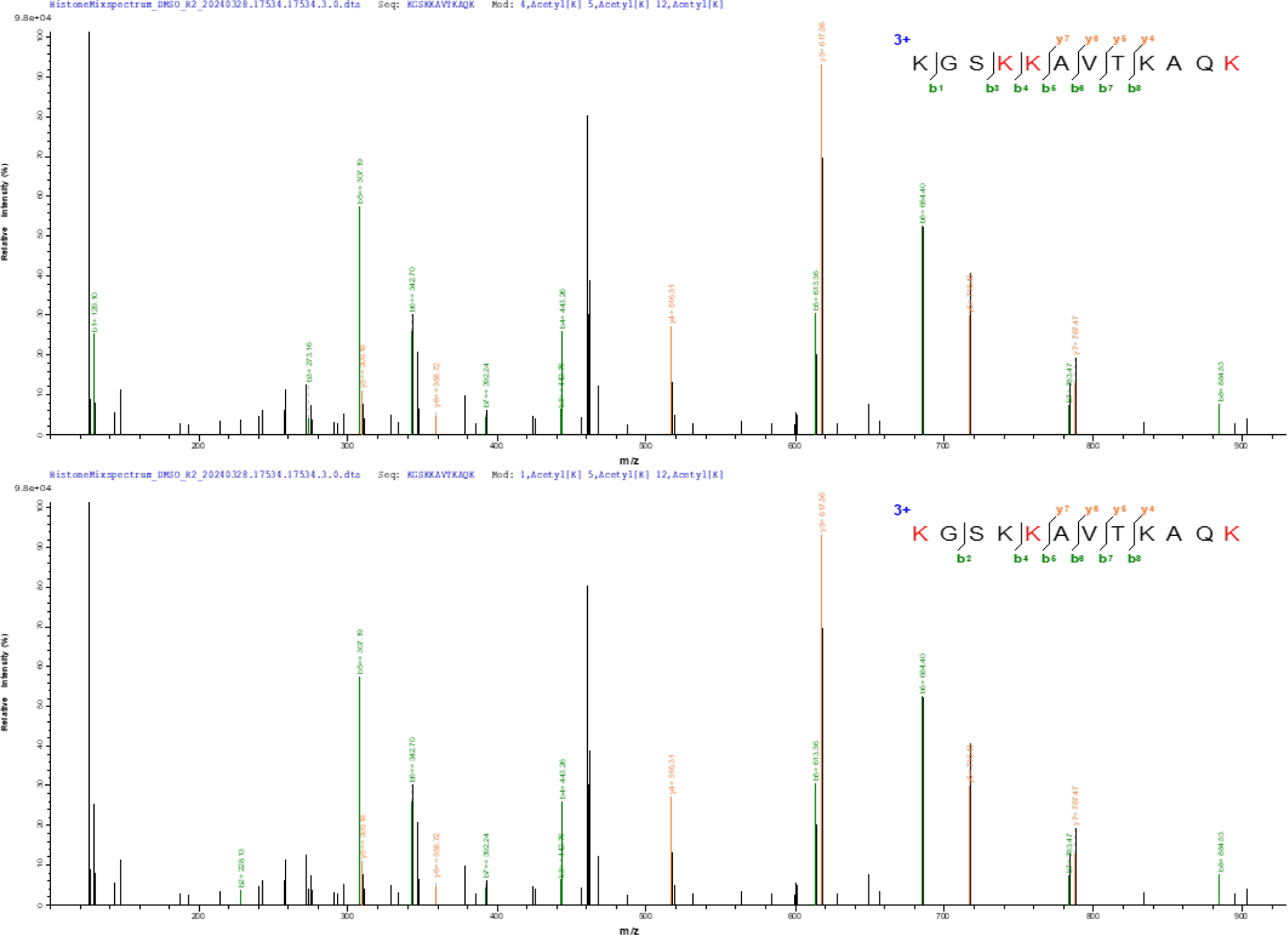
MS/MS spectral evidence supporting deconvolution in Fig. 5i.

**Supplementary Figure 9d.**
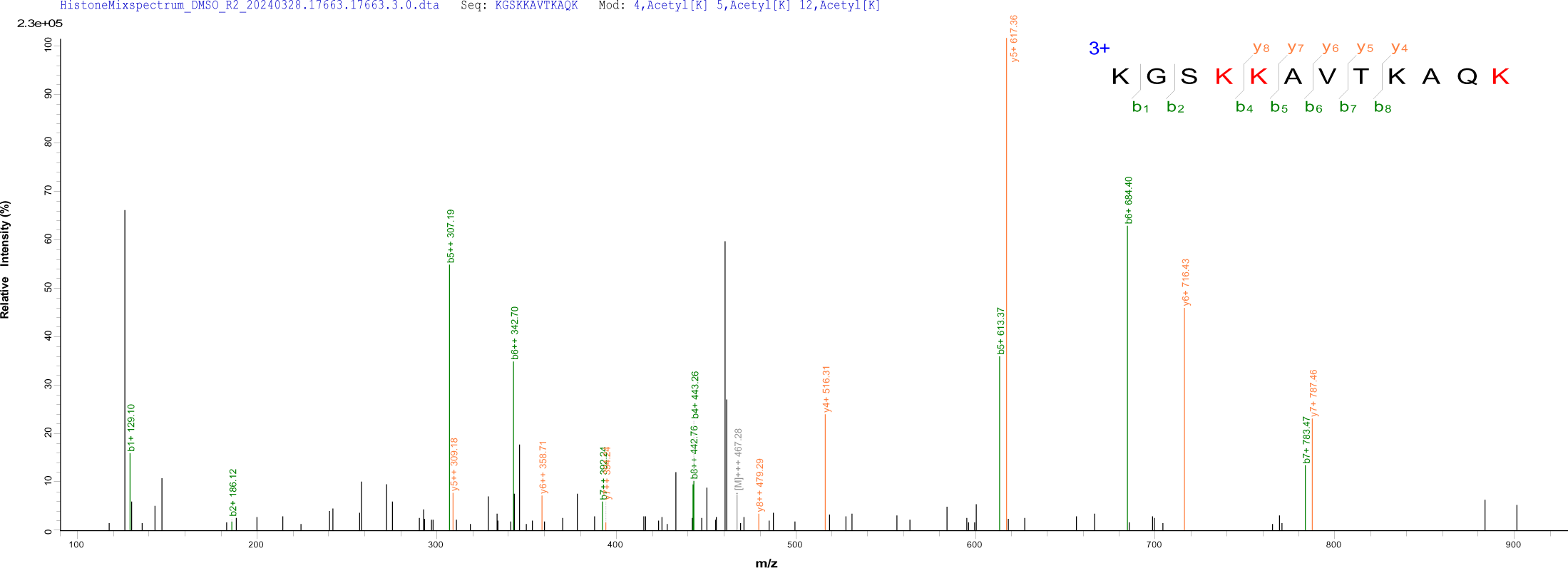
MS/MS spectral evidence supporting deconvolution in Fig. 5i.

**Supplementary Figure 10.**
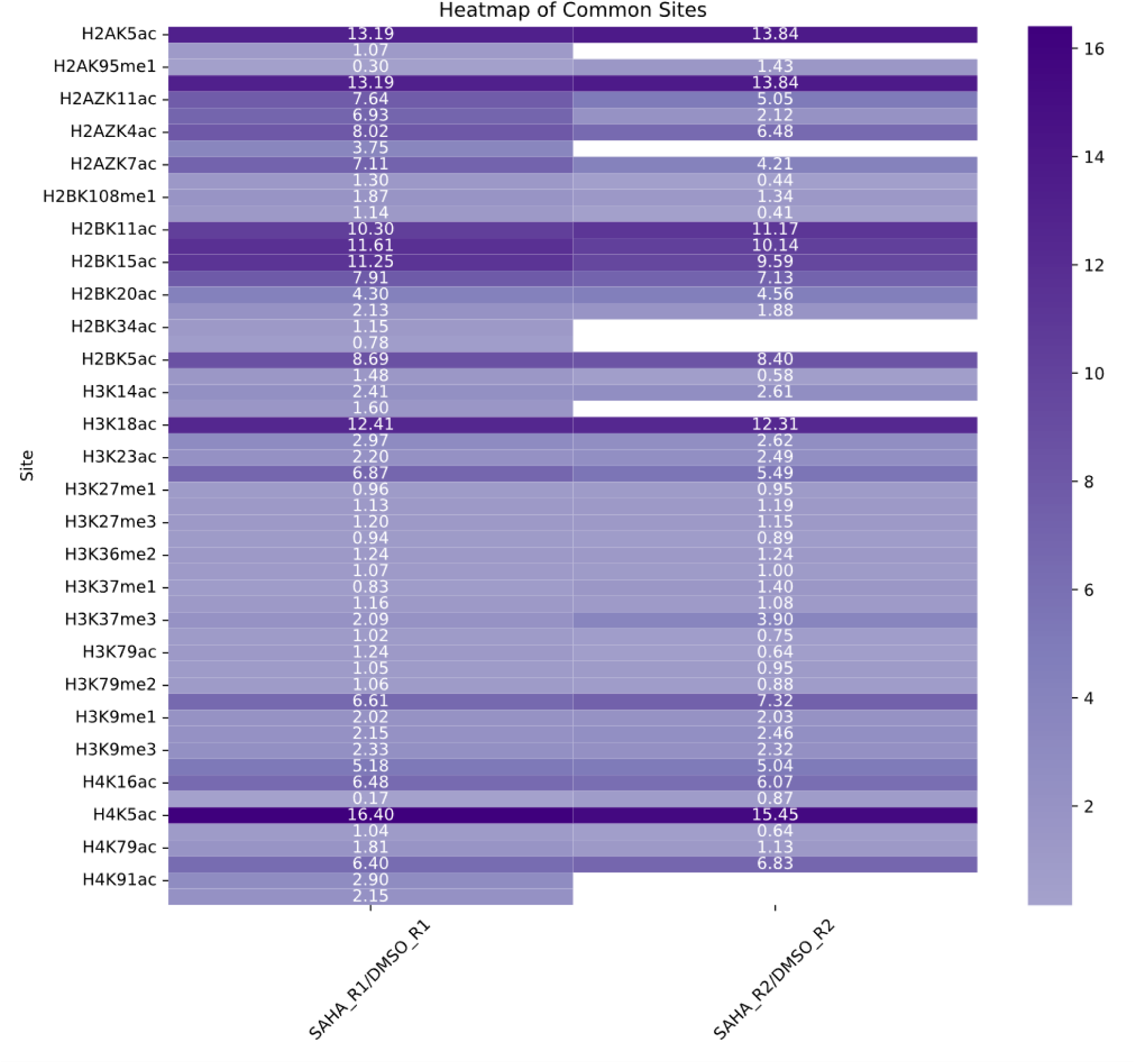
Reproducible PTM quantification by PTMdecoder. Highly correlated ratio of changes between two biological replicates (Fig. 5) for histone PTM sites identified by both standard PTMdecoder and top-1 mode workflows suggest consistent performance of PTMdecoder.

**Supplementary Figure 11.**
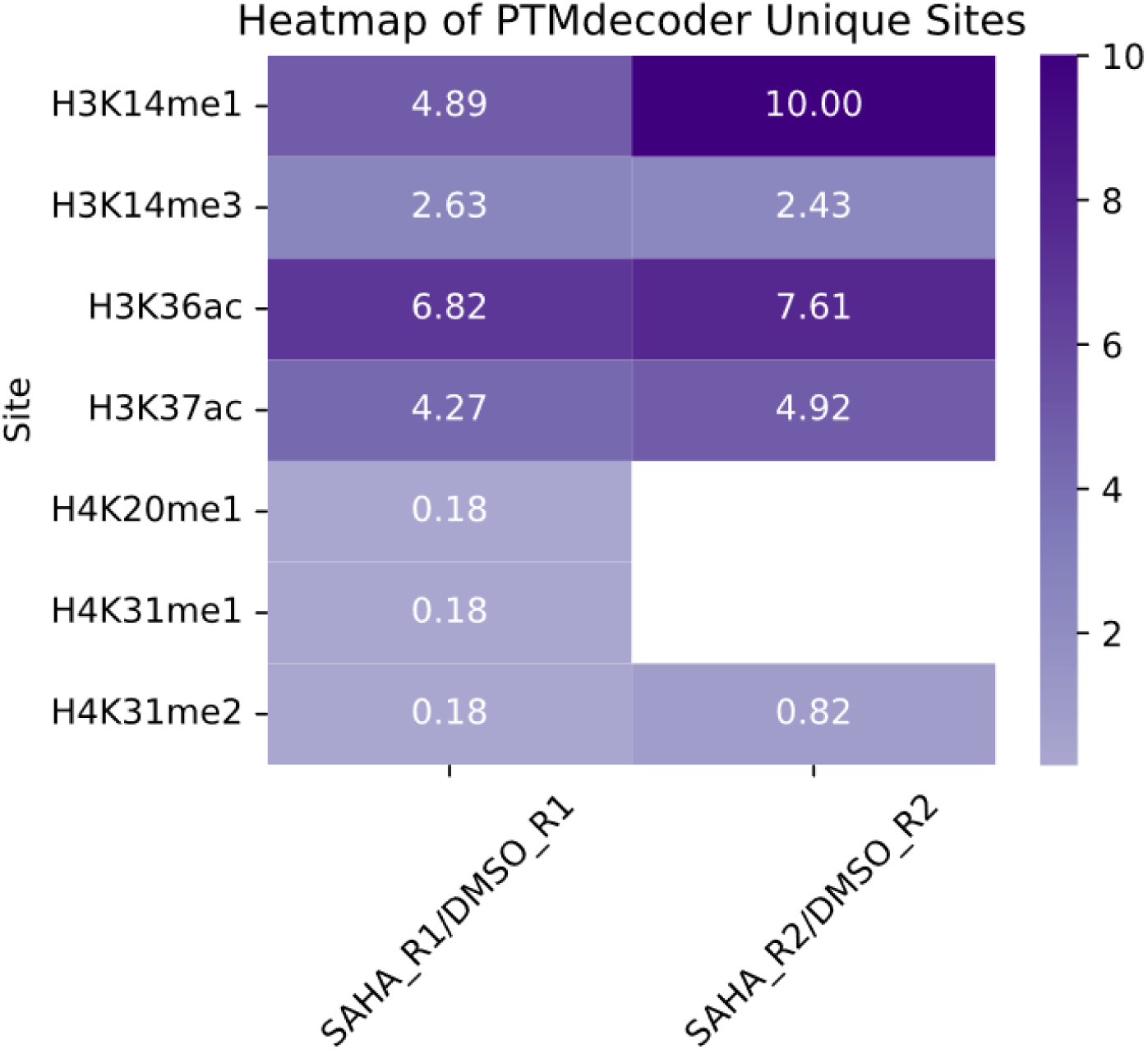
Expanded PTM site detection by PTMdecoder. Additional ratio of changes between two biological replicates (Fig. 5) for histone PTM sites uniquely identified by the standard PTMdecoder workflow indicate PTMdecoder’s capacity compared to the top-1 mode.

## Supplementary Notes

**Supplementary Note 1. Quantification based on XIC and proportions derived from MS/MS deconvolution**

1. XIC peak smoothing.

Intensities are smoothed along the retention time axis using a moving average filter. The original (unsmoothed) XIC is retained for subsequent filtering (Step 3).

2. Detection of XIC peaks for mixed IMPs.

Extracted ion chromatogram (XIC) peaks are detected and reconstructed based on MS1 spectra signals, centered around the identified MS/MS spectra. For each MS/MS-identified precursor, peak boundaries are defined at retention times where intensities fall below a user-defined threshold 𝛼 (default: 0.01) of the local maximum precursor intensities.

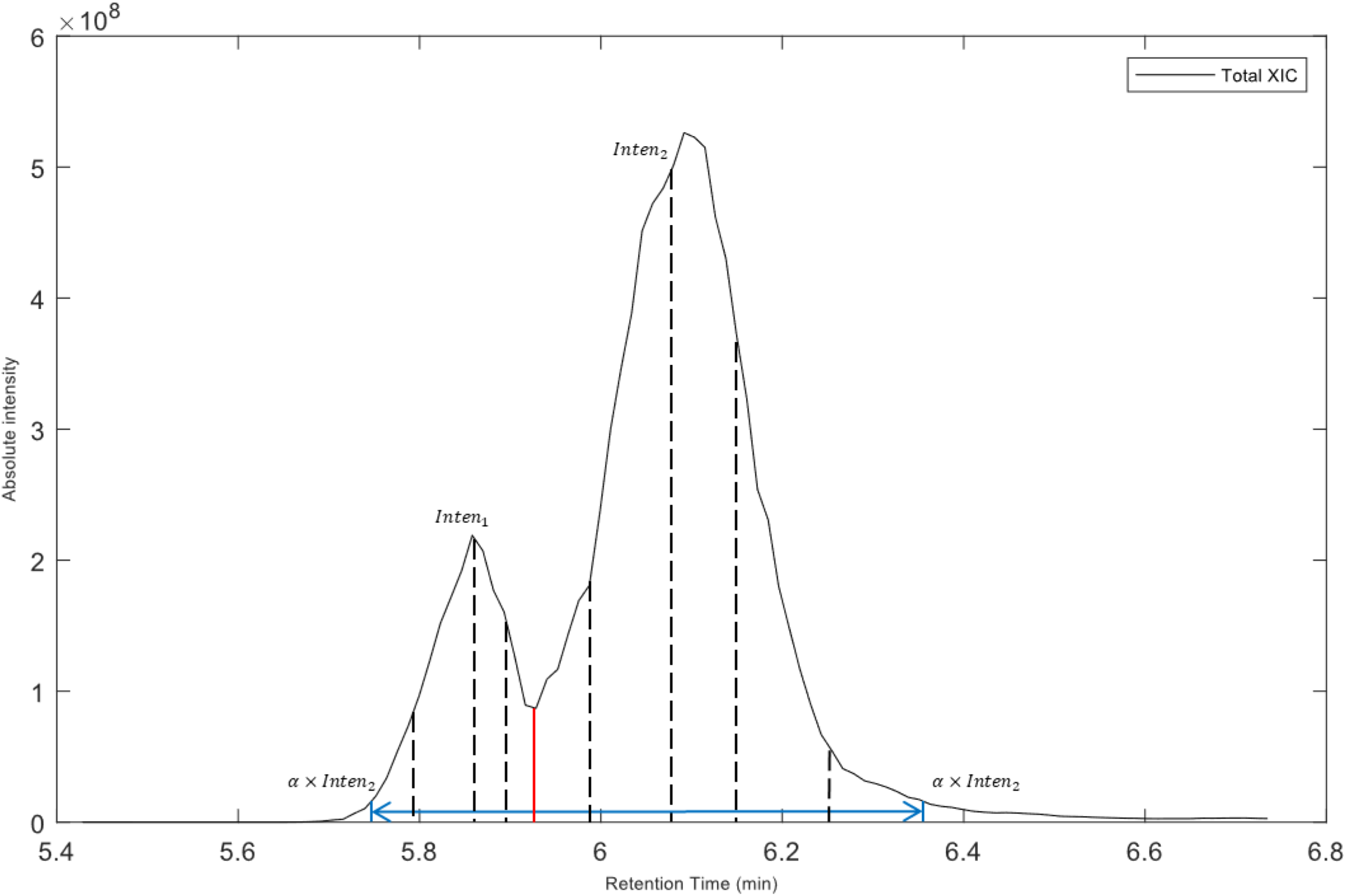

Peak boundaries (blue vertical lines in the figure) are defined at retention times where precursor intensities drop below 𝛼 × 𝐼𝑛𝑡𝑒𝑛_2_ — the smaller of the two local maxima (𝐼𝑛𝑡𝑒𝑛_2_ < 𝐼𝑛𝑡𝑒𝑛_1_). Peaks are truncated at local minima with intensities less than half of adjacent maxima on either side, splitting them into separate peaks.

3. Removing XIC peaks with insufficient MS1 spectrum support.

An XIC peak is removed if the corresponding original XIC peak is supported by fewer than 5 matched MS1 peaks. This step mitigates false positives: narrow peaks may reflect noise, even if peptide-spectrum matches (PSMs) are present.

4. MS/MS-guided XIC deconvolution.

The relative abundance profiles of individual peptidoforms are estimated through a non-parametric approach. For each retention time point 𝑡 within a XIC peak for mixed IMPs, the proportion of an IMP was calculated by applying Nadaraya-Watson kernel regression to MS/MS-derived proportions from nearby elution times. Specifically, the estimated proportion 𝑟^(𝑡) was computed as:

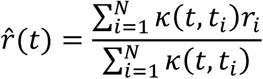

where 𝑟_𝑖_ are the proportion determined by MS/MS deconvolution at time 𝑡_𝑖_, and 𝜅(𝑡, 𝑡_𝑖_) is the kernel weighting function (Gaussian kernel used here since it has an untruncated property). The kernel bandwidth is determined according to Silverman’s rule of thumb:

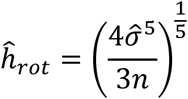

where 𝑛 is the number of identified MS/MS spectra in the retention time range of this XIC peak, and 𝜎^ is the standard deviation of the retention times for these identified MS/MS precursors. Then, the proportions at each retention time are normalized to ensure that the sum of the proportions is equal to 1.

5. Low-abundance IMP elimination

To eliminate the false positives, the deconvoluted XIC peaks of peptidoforms are considered unreliable and removed if the areas under the peaks are less than 10% of the highest-abundant co-eluting peptidoform. The areas of these excluded peptidoforms are redistributed to remaining peptidoforms using the method above.

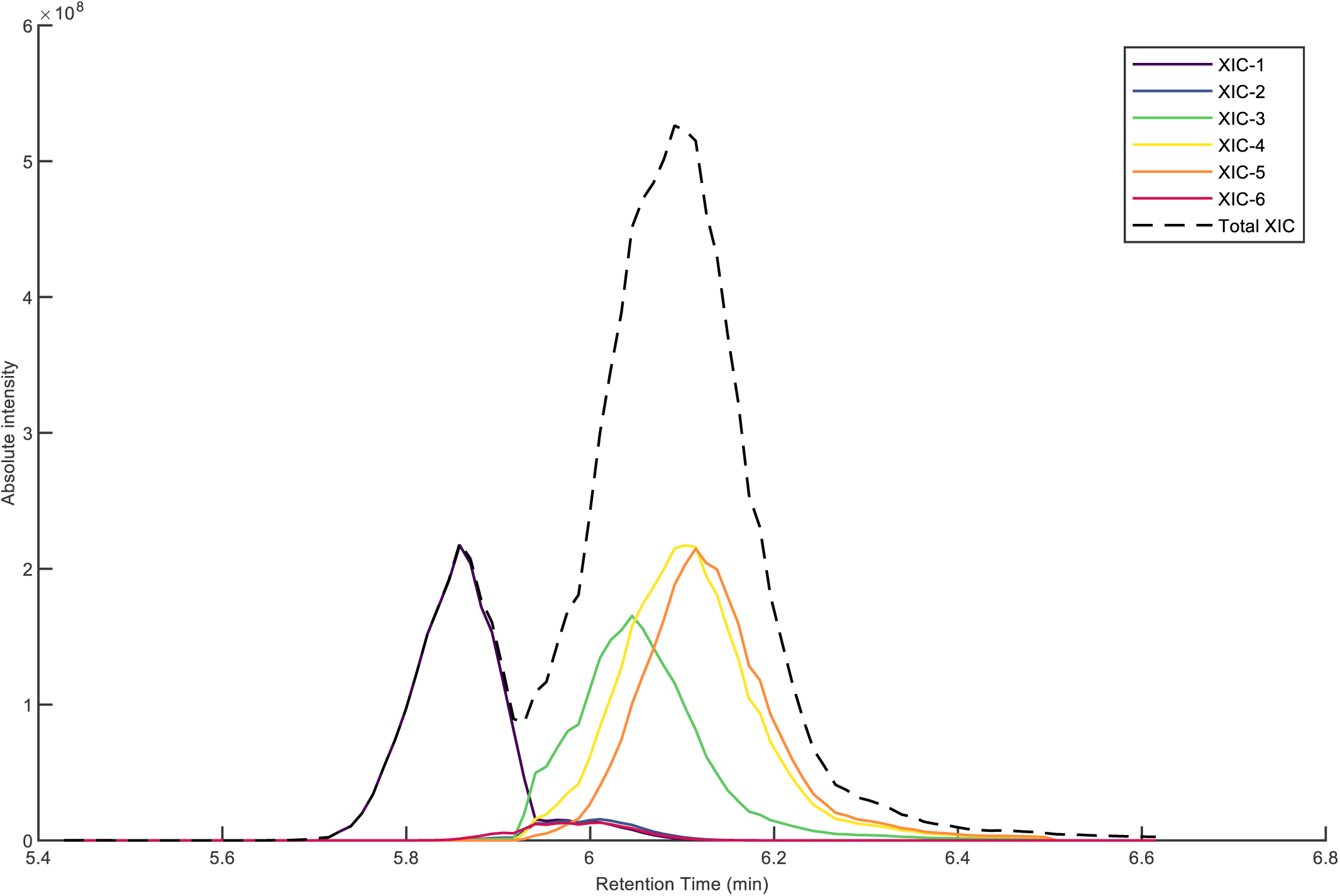

The above figure shows an example of XIC peak elimination. In the right peak, ‘XIC-2’ and ‘XIC-6’ (areas <10% of ‘XIC-4’) are eliminated, and their areas are reassigned to other peptidoforms, as shown in the following figure.

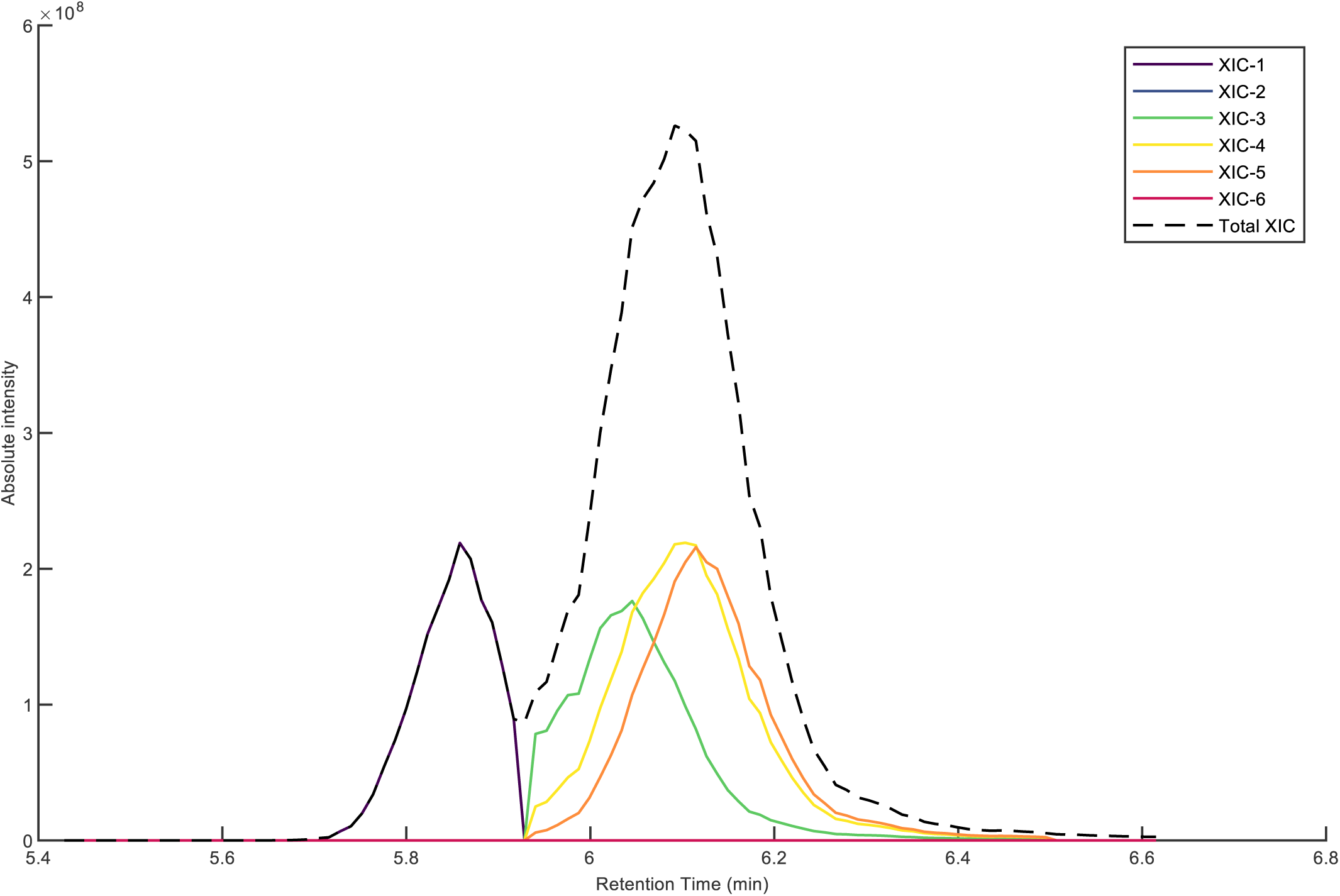

6. Selection of a Single XIC Peak per Peptidoform.

Each peptidoform should be quantified by exactly one XIC peak. If multiple candidate peaks exist (e.g., due to misidentified MS/MS spectra), only the peak with the largest full width at half maximum (FWHM) is retained, as indicated by the blue arrows in figure below.

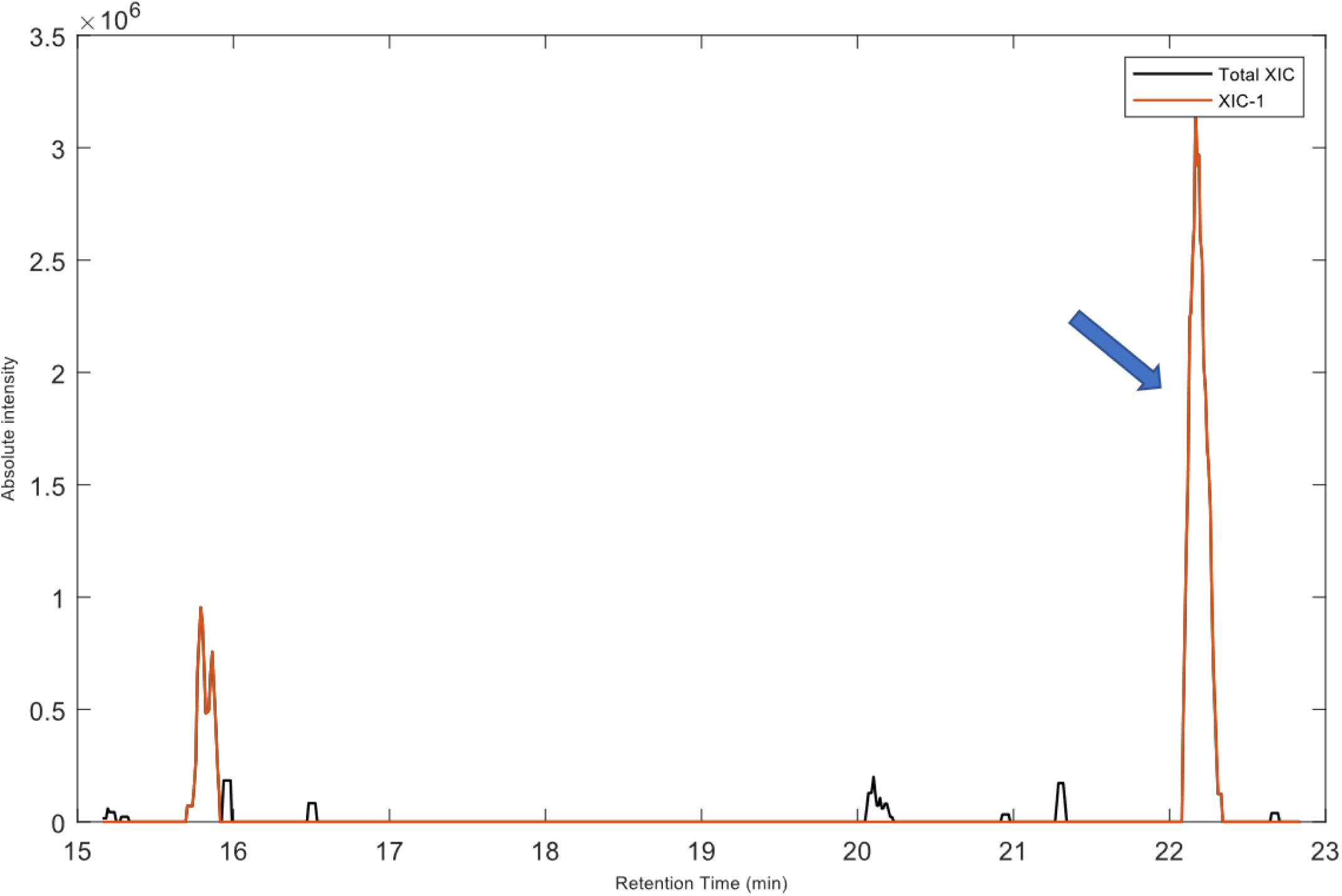

7. Quantification via Area Integration

The abundance of each peptidoform is calculated as the integrated area under its reconstructed XIC curve.

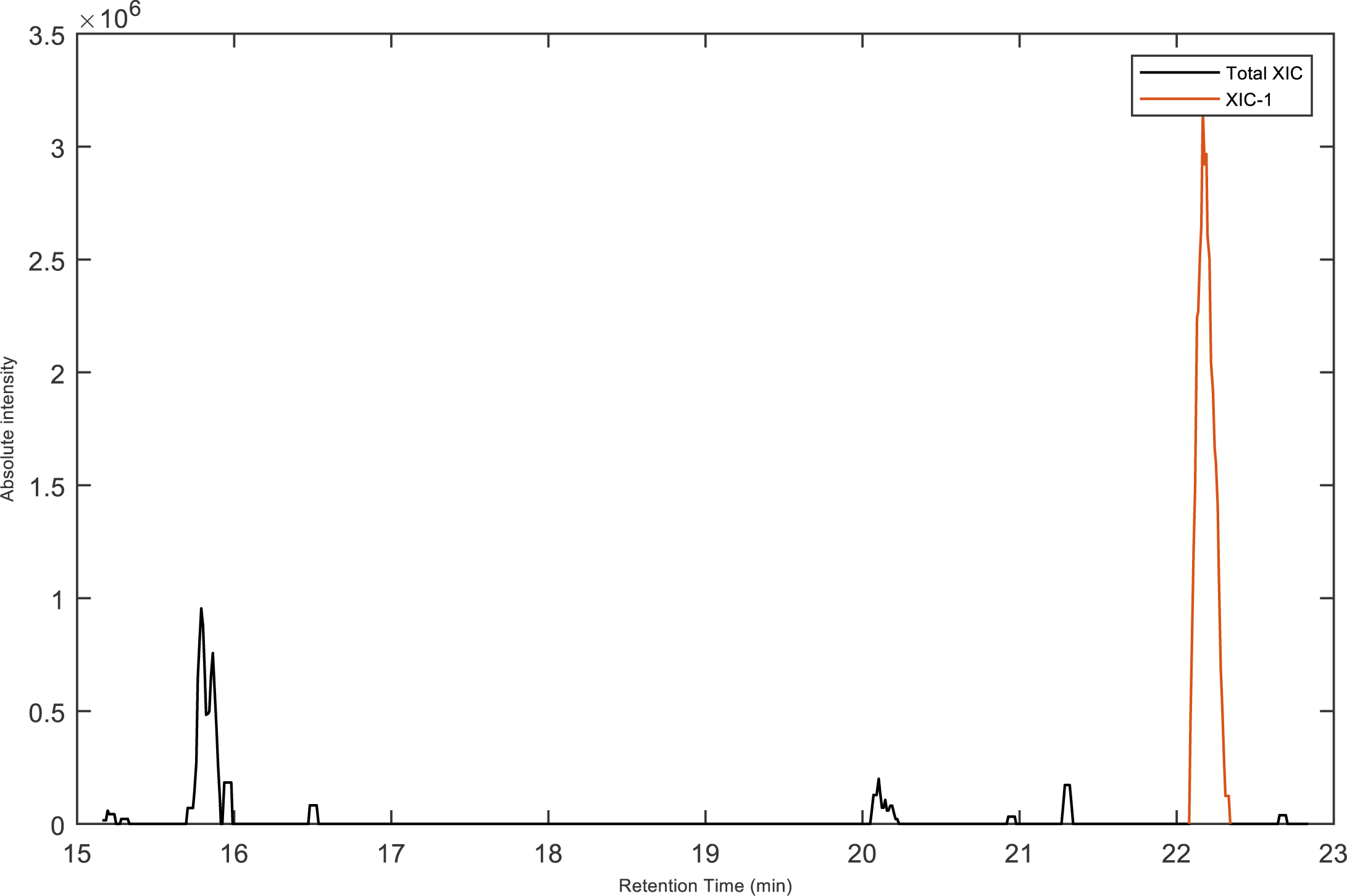

## Notes

### Competing Interest Statement

The authors have declared no competing interest.

